# Gene-Expression Programs in Salivary Gland Adenoid Cystic Carcinoma Analyzed Using Single-Cell and Spatial Transcriptomics

**DOI:** 10.1101/2025.09.01.673548

**Authors:** Ifeoma Ebinumoliseh, Gopikrishnan Bijukumar, Kendall Hoff, Kathryn J. Brayer, Elaine L. Bearer, Scott A. Ness, Jeremy S. Edwards

**Author notes:** Contributed equally (Ifeoma Ebinumoliseh, Gopikrishnan Bijukumar).

## Abstract

Adenoid cystic carcinoma of the salivary gland (SGACC) is a highly aggressive malignancy characterized by poor patient survival outcomes. While several studies have analyzed the transcriptome of the salivary gland at the bulk and single-cell level, no spatial transcriptomic analyses of this tissue have been published. Most of the existing publications on SGACC have predominantly relied on bulk and single cell RNA sequencing approaches, which do not resolve the spatially localized transcriptional heterogeneity nor have the resolution for defining molecular markers within tumor subpopulations. SGACC is clinically notable for the presence of multiple tumor clones, distinct spatial phenotypes, and its indolent yet invasive nature coupled with a high propensity for distant metastasis. These features may reflect co-expression of tumor-associated markers across diverse cellular niches, and a resultant biological complexity which causes standard treatment such as surgical resection, radiation therapy, and chemotherapy to be largely ineffective in significantly improving long-term survival, and highlights the need for more precise, targeted therapeutic strategies. Herein, we analyzed single cell (n = 4) and high-resolution spatial transcriptomics samples (n = 5) to characterize cancer cell populations in MYB- and non-MYB-expressing cell states, delineated gene expression signatures, and identified critical molecular interactions specific to SGACC. We used Visum HD to obtain spatial transcriptomics data at 2µm squared high resolution. This allowed a multi-omics approach comprising single cell and spatial transcriptomic methods to enable the discovery of novel transcriptional signatures and microenvironmental features not captured by conventional methods. Spatial mapping revealed marked cellular heterogeneity and demonstrated how tissue environments influence cellular transcriptomics. To tumor heterogeneity, we focused on tumorigenic cell populations, profiled plasma and T cell enrichment within the tumor microenvironment and identified key pathways and transcriptional drivers including the MYB-NFIB fusion underlying the tumor cluster formation. Our findings indicate an upregulation of genes involved in extracellular matrix remodeling, autophagy, and reactive stromal cell populations. We further found evidence of partial epithelial-mesenchymal transition (P-EMT) programming within MYB-expressing tumor clusters. Pathway analysis revealed that mutations in the spatial query sample prominently affect the PI3K-AKT and IL-17 signaling pathways, together with a downregulation of canonical Wnt signaling in some regions of the tissue architecture adjacent to immune cells. Collectively, these results underscore the complex regulatory landscape of SGACC and offer insights into its cellular dynamics and possible therapeutic vulnerabilities.

## 1. INTRODUCTION

Salivary gland adenoid cystic carcinoma (SGACC) is an aggressive malignancy and ranks among the most prevalent neoplasms of the salivary glands^1^. It is characterized by slow, yet locally invasive growth, and a high propensity for perineural invasion and distant metastasis. These features often result in the colocalization of diverse transcriptional profiles and subsequent clonal evolution^1,2^. Conventional treatment modalities including surgical resection, radiation therapy, and chemotherapy, lead to highly variable patient outcome with a reported median survival range from 16.8 to 120 months, and a post-operative survival extending from a few months to under 15 years^1^. Although salivary gland malignancies often progress asymptomatically and are generally indolent, they may result in functional neurological deficits and are associated with poor long-term prognoses^3–5^.

SGACC typically arises in one of the three major craniofacial secretory glands which are the parotid, submandibular, and sublingual glands as well as in minor salivary glands, which collectively account for 70–90% of salivary gland malignancies^6,7^. The tumorigenic process in SGACC is driven by specific genomic alterations, notably chromosomal rearrangements such as t(6;9), 6q22–23 and 8q13 translocations, which lead to fusion events or aberrant activation of MYB, MYBL1, and NFIB genes. These alterations impact several key oncogenic pathways, including NOTCH, PI3KCA, and PTEN ^8–12^. The prevalence of the MYB–NFIB fusion gene varies substantially across cohorts from 16% to nearly 100% owing to variability in the fusion breakpoints. These fusions typically involve MYB exon 14 and NFIB exon 9, producing either full-length or truncated MYB fusion transcripts. Despite truncation, these variants retain functional DNA-binding and transactivation domains, underscoring the pivotal role of MYB in transcriptional regulation within SGACC ^9,13–15^.

While prior bulk RNA sequencing analyses of ACC have identified important biomarkers, they have been unable to resolve tumor-specific transcriptional profiles from individual phenotypes limiting insights into cell-specific tumor biology. SGACC poses additional complexity due to its biphasic architecture, which arises from the presence of both ductal and myoepithelial components^16^, along with the co-localization of epithelial and mesenchymal cell populations^17^. This mixture of cells contributes to increased metastatic potential, and persistent therapeutic resistance^18^.

Given the limitations of bulk RNA sequencing in collectively resolving the tumor architecture, the next step is to employ a multi-omics approach including single cell and high-resolution spatial transcriptomics to assess transcriptional colocalization and delineate tumor clonal framework. In this study, we integrated single-cell and spatial transcriptomics to characterize key transcriptional signatures and biological pathways enriched within SGACC cell sub-populations. This high-resolution approach enabled a detailed investigation of cancer cell plasticity and intercellular crosstalk between and within MYB and non-MYB expressing cell states. Utilizing Cell2Location for cellular deconvolution and visualization, our analysis revealed interactions between immune cells such as Plasma cells, T-lymphocytes and core tumor cell clusters. These interactions were supported by distinct differentially expressed gene signatures associated with tumor progression, loss of cell-cell adhesion, and a partial retention of adhesion markers paired with an aberrant negative regulation of canonical Wnt signaling in certain subsection of the tissue. This was found to be in concert with other molecular fingerprints which suggested the presence of partial epithelial-to-mesenchymal transition (P-EMT) within the tumor. Furthermore, we uncovered normal cell expression in cells spatially positioned at the periphery of MYB-expressing regions in the tissue which may be residual normal non-malignant cell. Additional findings underscore that many of the defining tumorigenic features in SGACC may be driven by MYB-expressing cell populations. Furthermore, this spatially resolved analysis highlights novel candidate pathways and molecular targets with potential therapeutic relevance.

## 2. Materials and Methods

### 2.1 Single cell samples

We analyzed a total of four distinct single-cell RNA sequencing (scRNA-seq) datasets in the published literature in this study (*n* = 4), comprising two normal salivary gland samples^21,22^ and two SGACC samples^19,20^. Three of the four datasets served as a reference for training the cell2location model for spatial mapping of cell populations. The normal salivary gland samples (N1 and N2) additionally served as controls in the differential expression analysis performed using PyDESeq2. Integration of three single-cell datasets was carried out using the integration function in Seurat, which was subsequently used to generate the reference UMAP (Fig. 1C).

**Figure 1.**
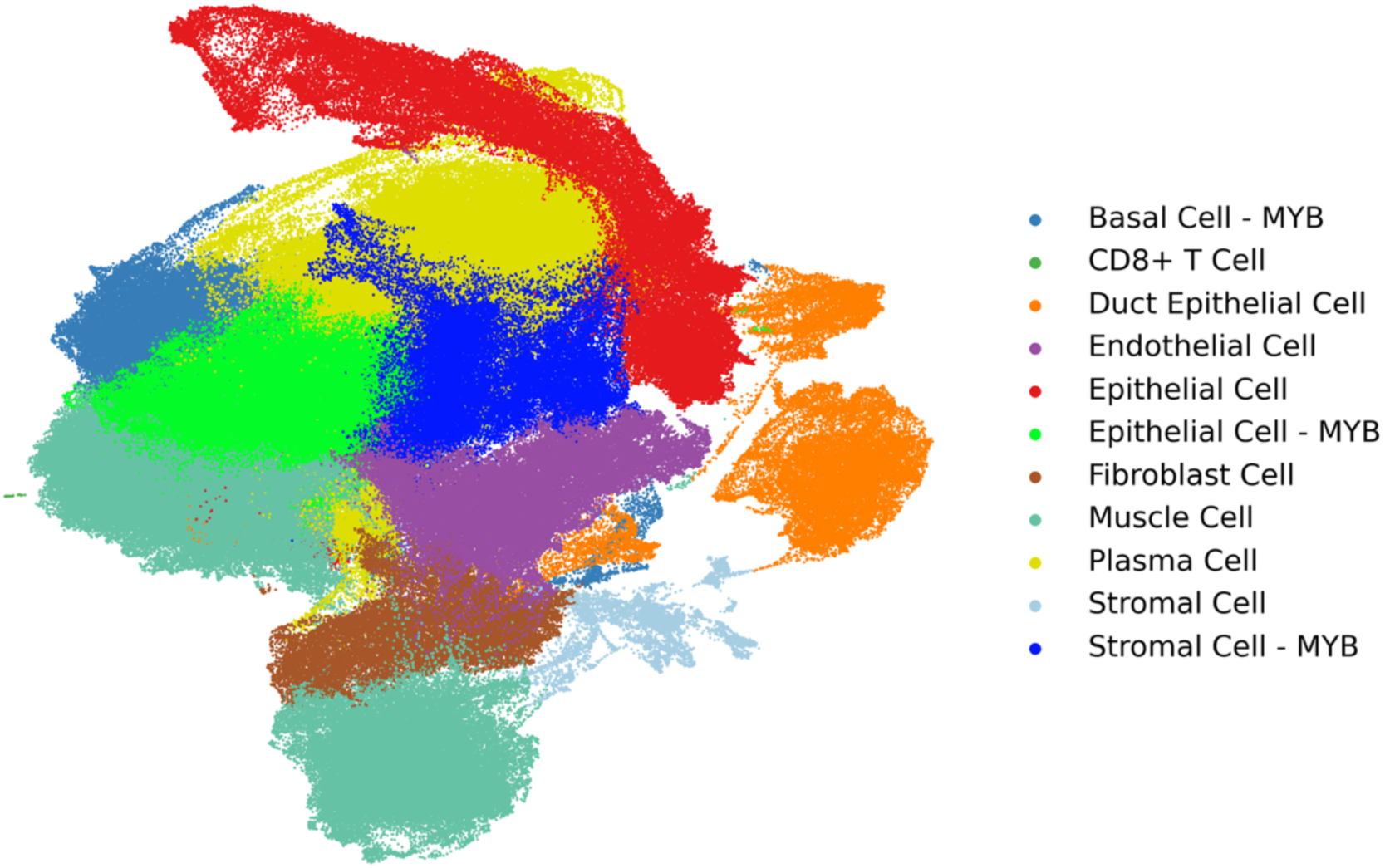
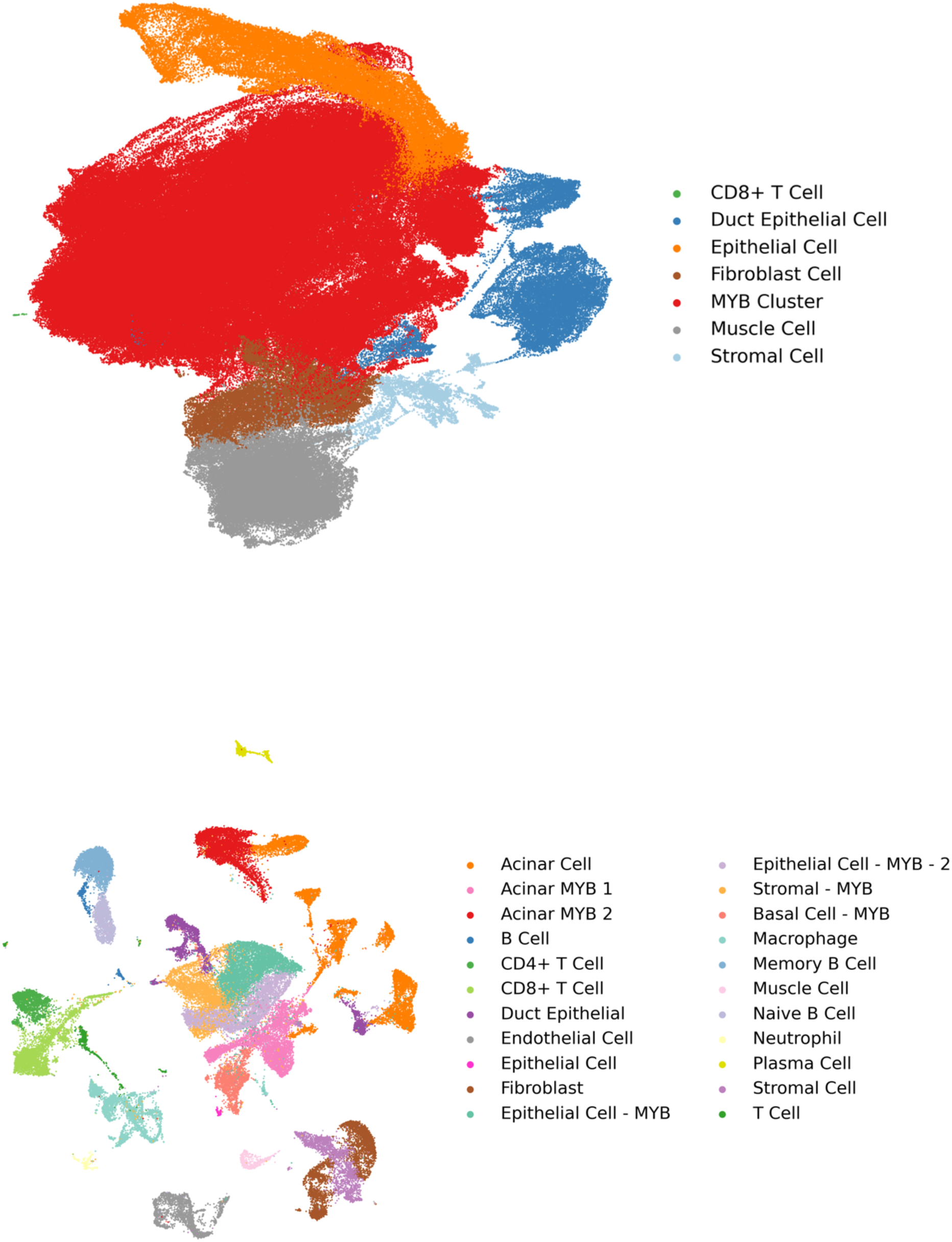
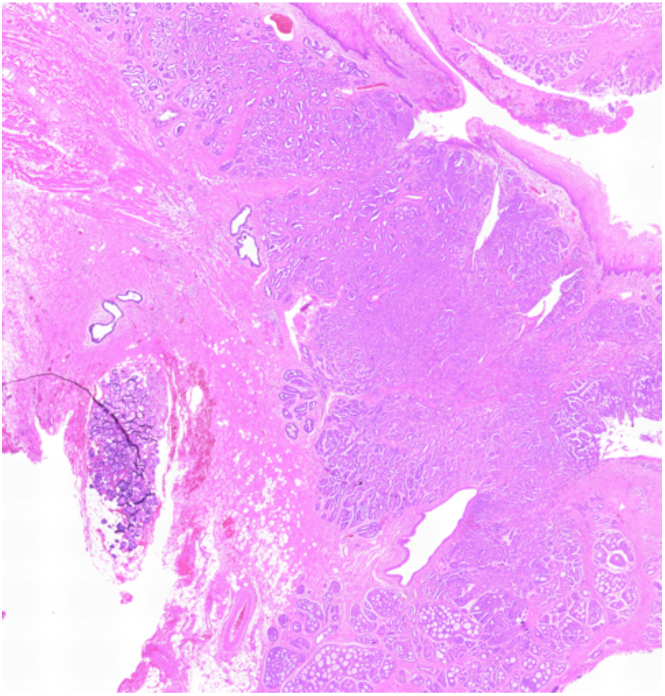
Identification and classification of cell populations using UMAP and histology. (A) UMAP visualization of cell clusters in the spatial query dataset. (B)UMAP highlighting MYB-expressing versus non-MYB-expressing cell clusters. (C). UMAP of integrated single-cell reference datasets. (D) H&E image of the SGACC spatial transcriptomics sample (ACC1)

### 2.2 Spatial Transcriptomics Samples

We analyzed five spatial SGACC samples (ACC1–ACC5; *n* = 5). ACC1 served as the primary sample for spatial omics investigation, while all five samples (ACC1–ACC5) were used collectively for differential expression analysis using PyDESeq2, with ACC2–ACC5 serving as biological replicates of SGACC (Fig. 2A-2B). The differentially expressed genes in PyDESeq2 analysis can be found in supplementary information 2 and 3. For each individual specimen, Formalin-Fixed, Paraffin-Embedded (FFPE) human salivary gland samples were used and processed according to the 10X Genomics Visium HD protocol CG000520 Rev B, with reagents provided by 10x Genomics^23,24^.

**Figure 2.**
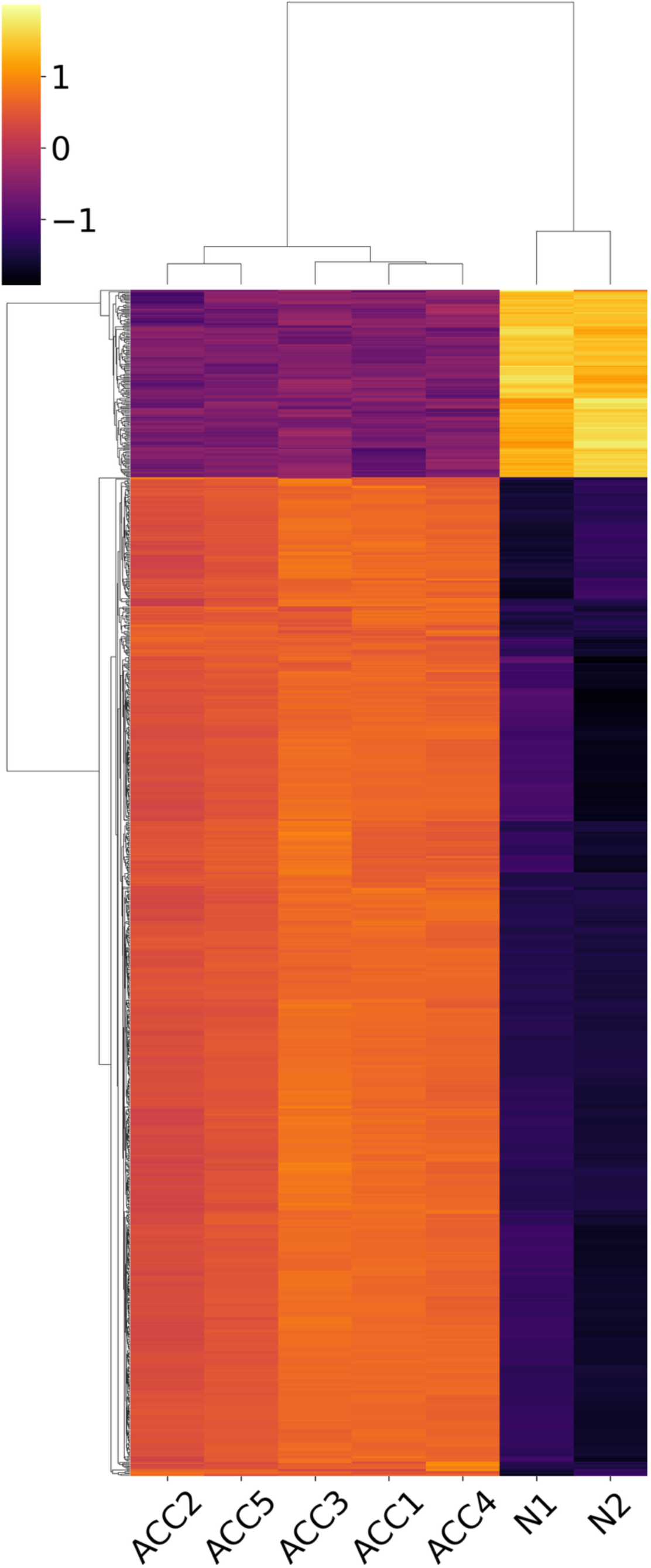
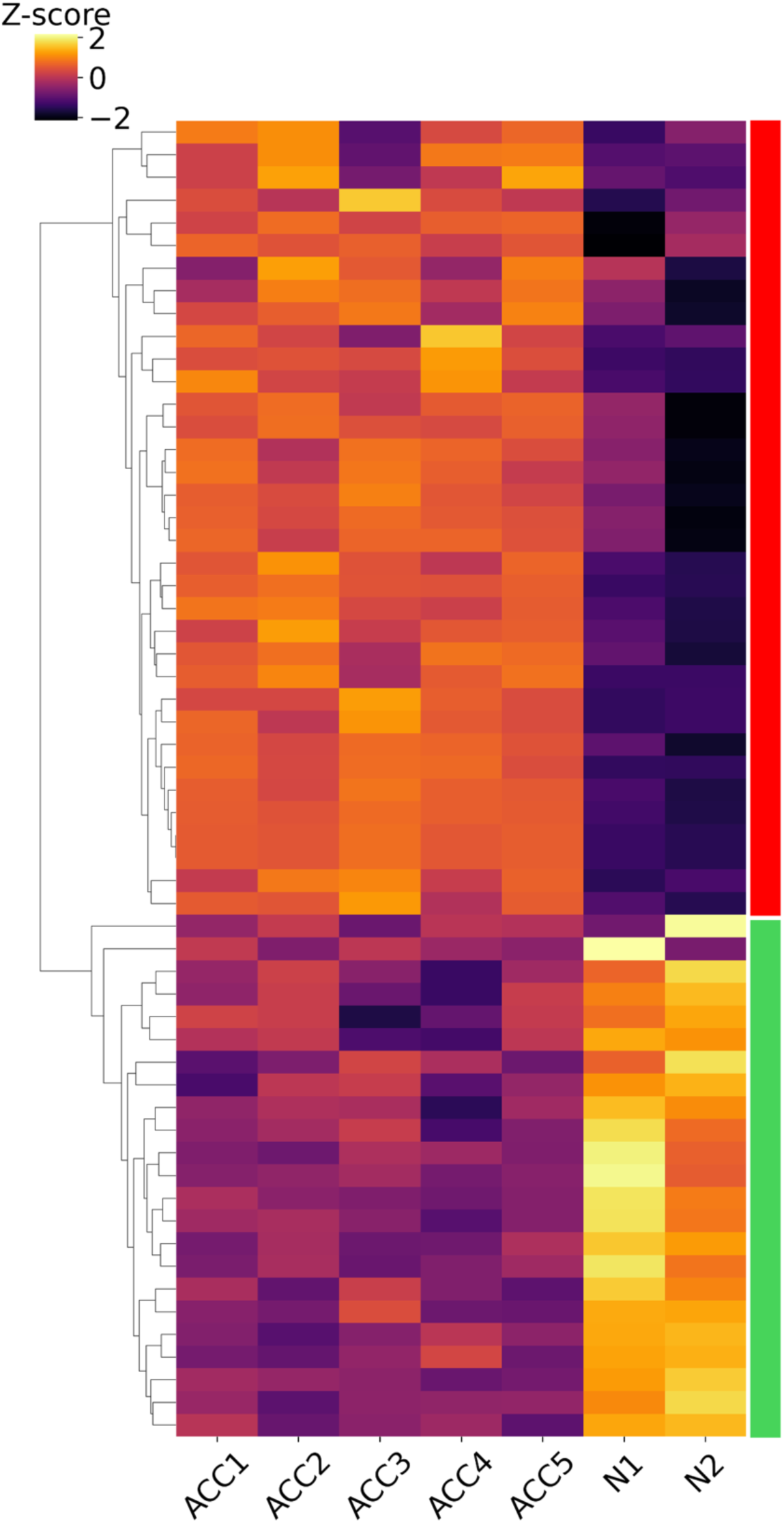
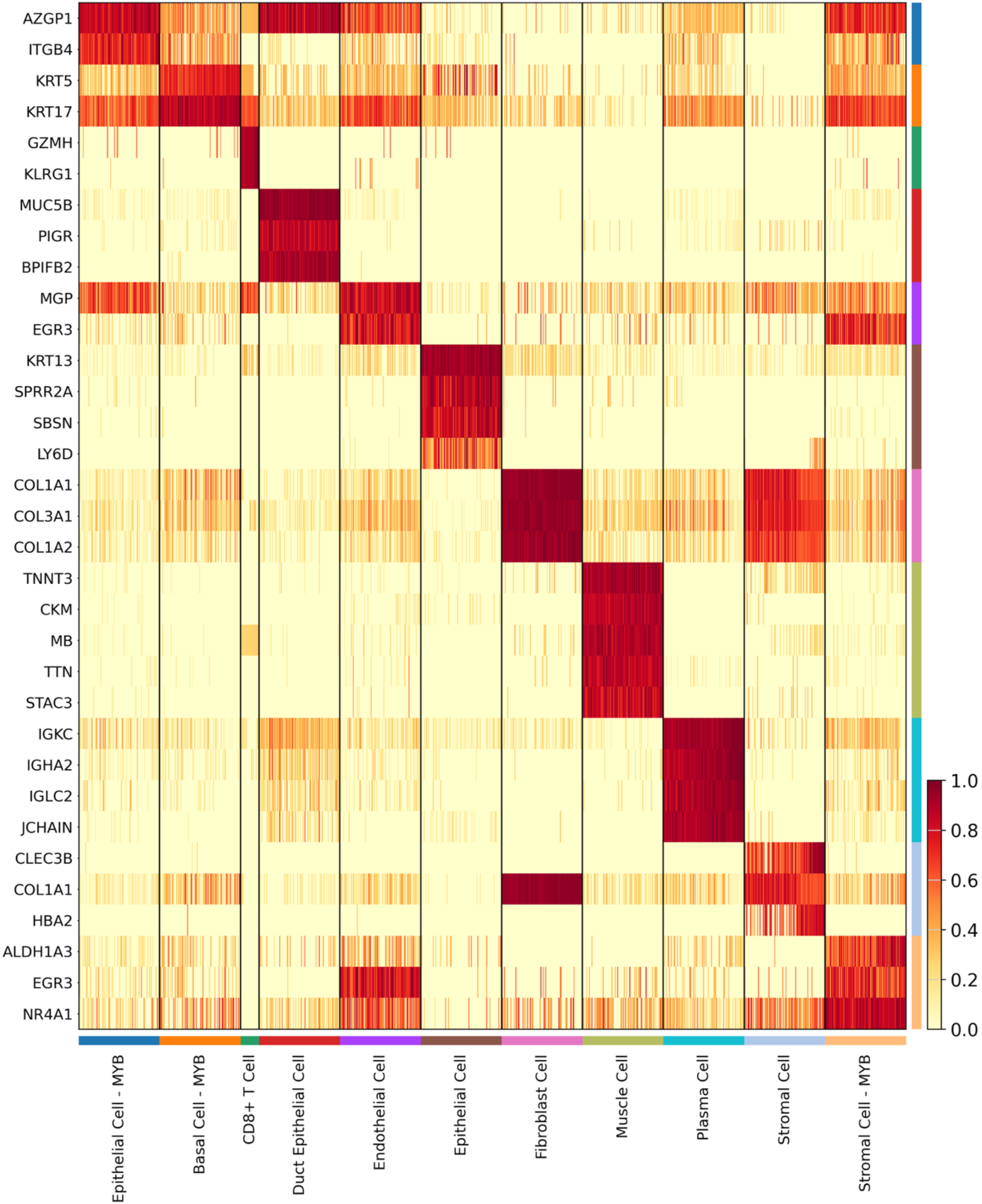
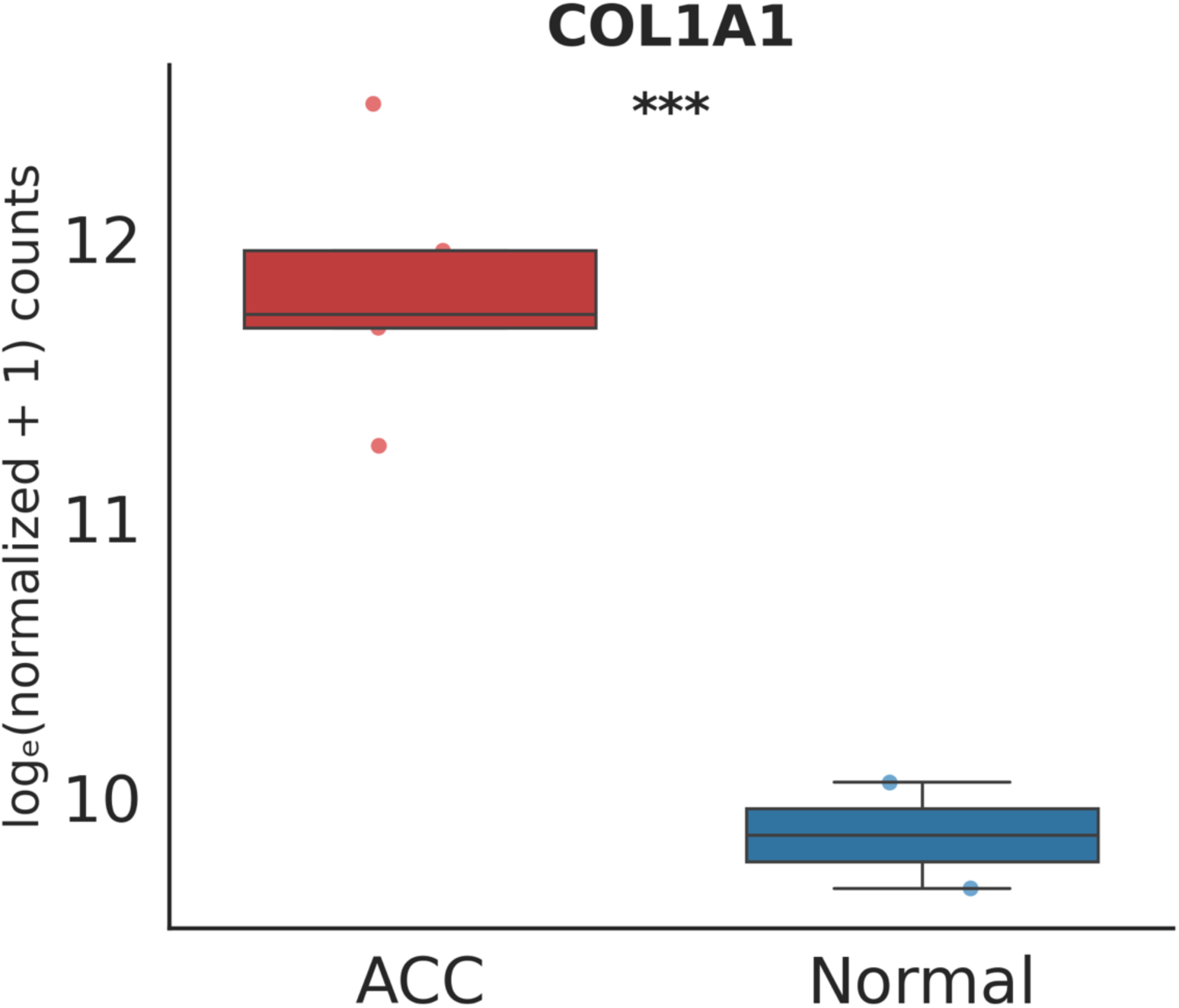
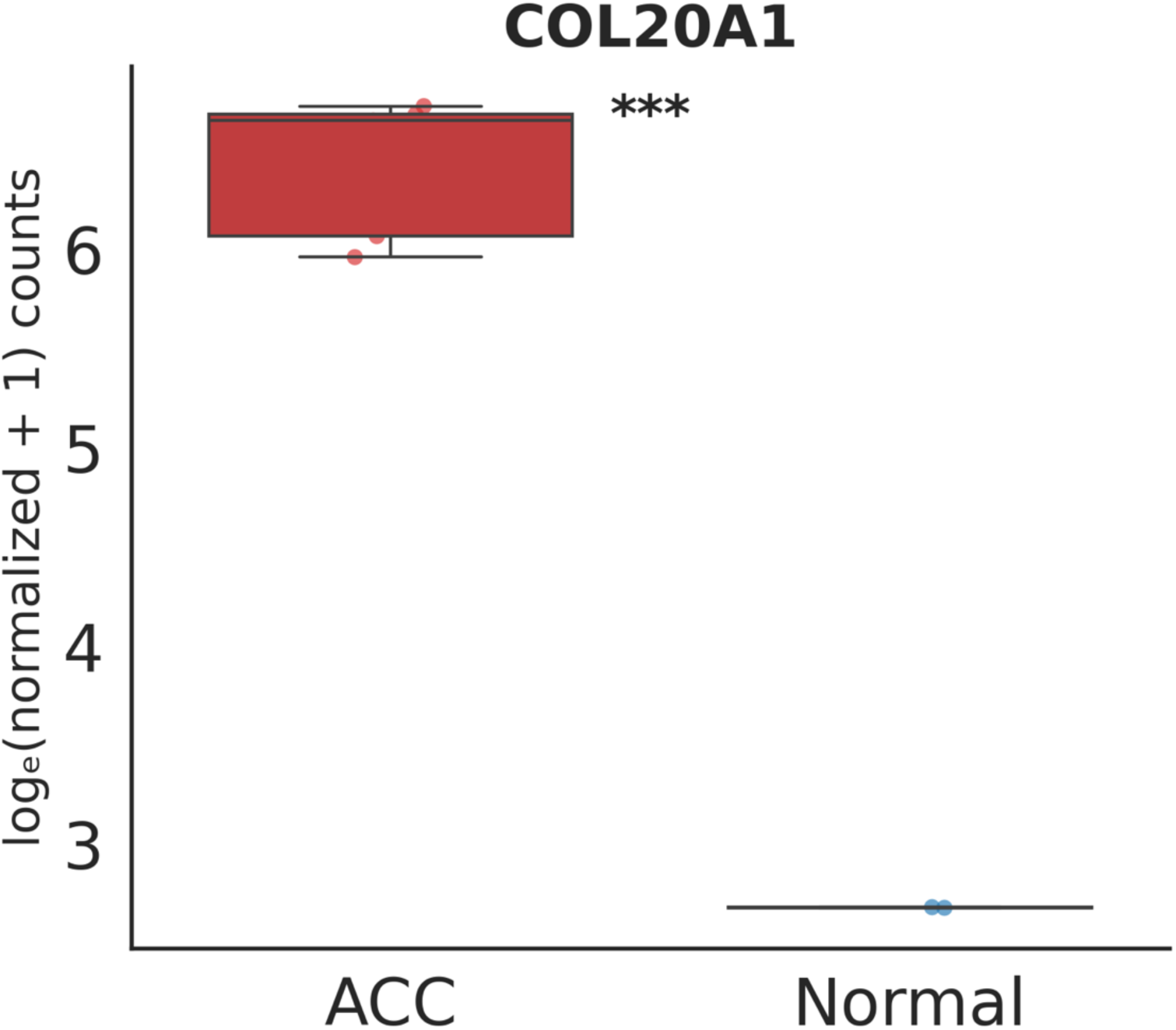
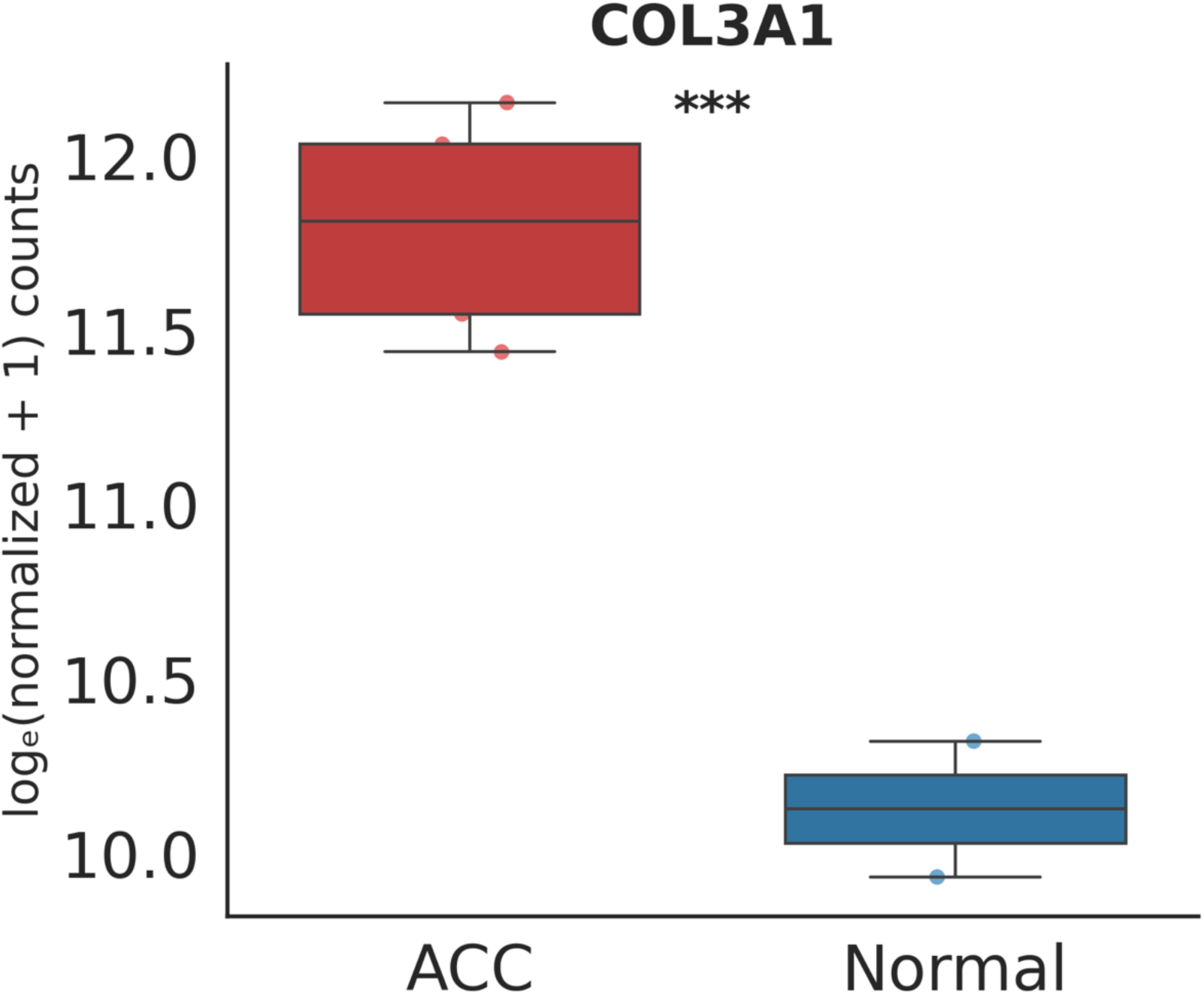
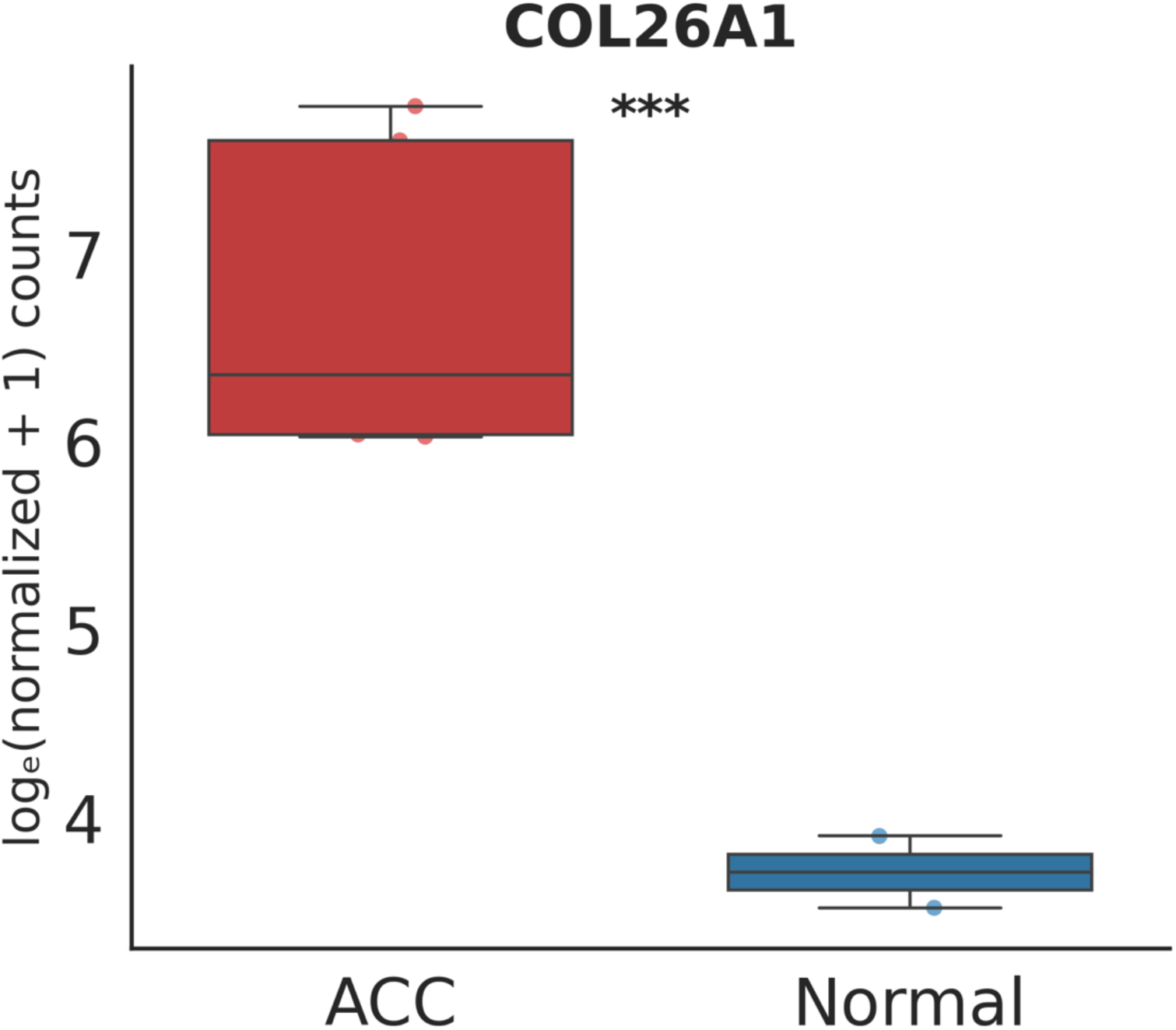
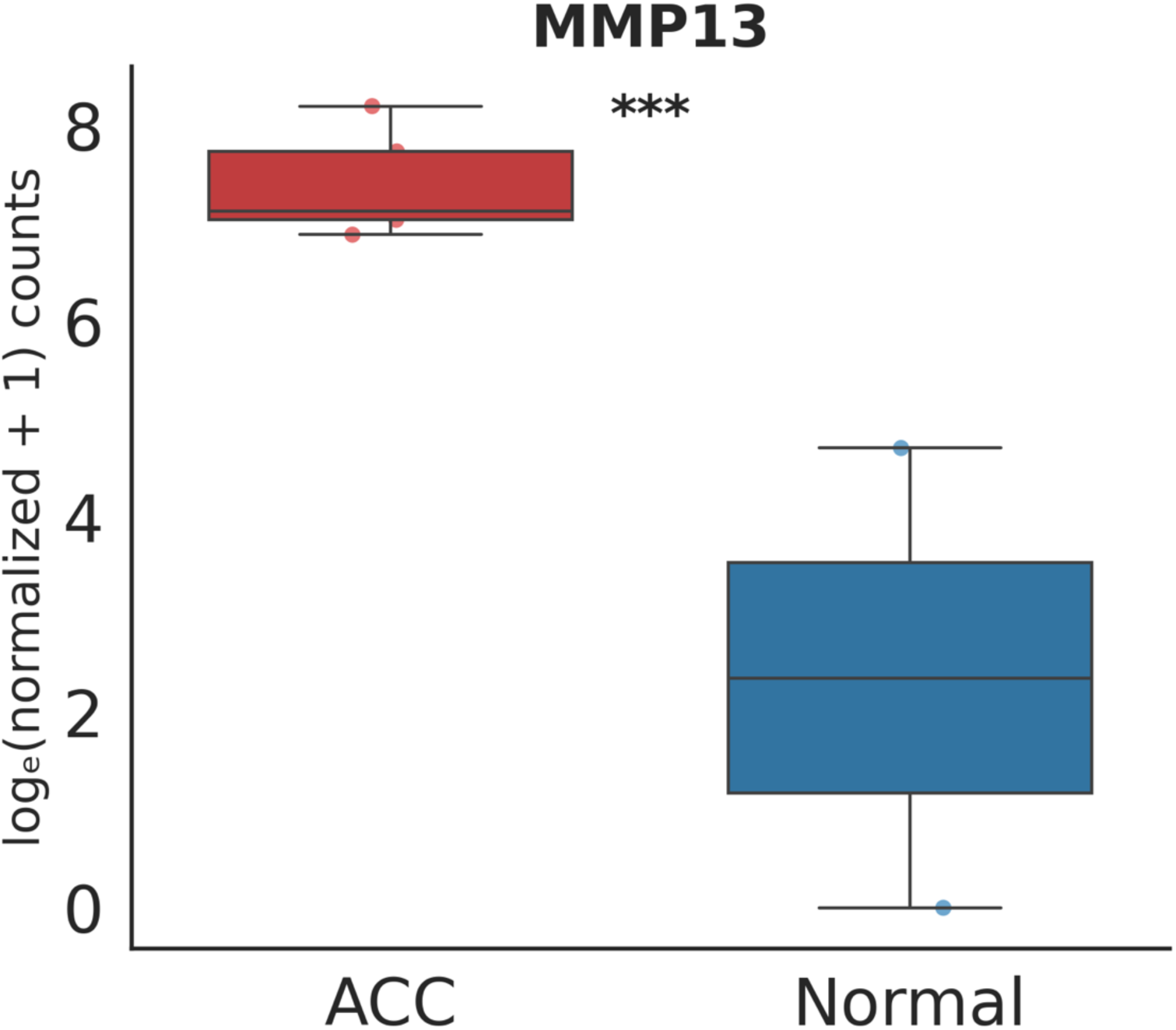
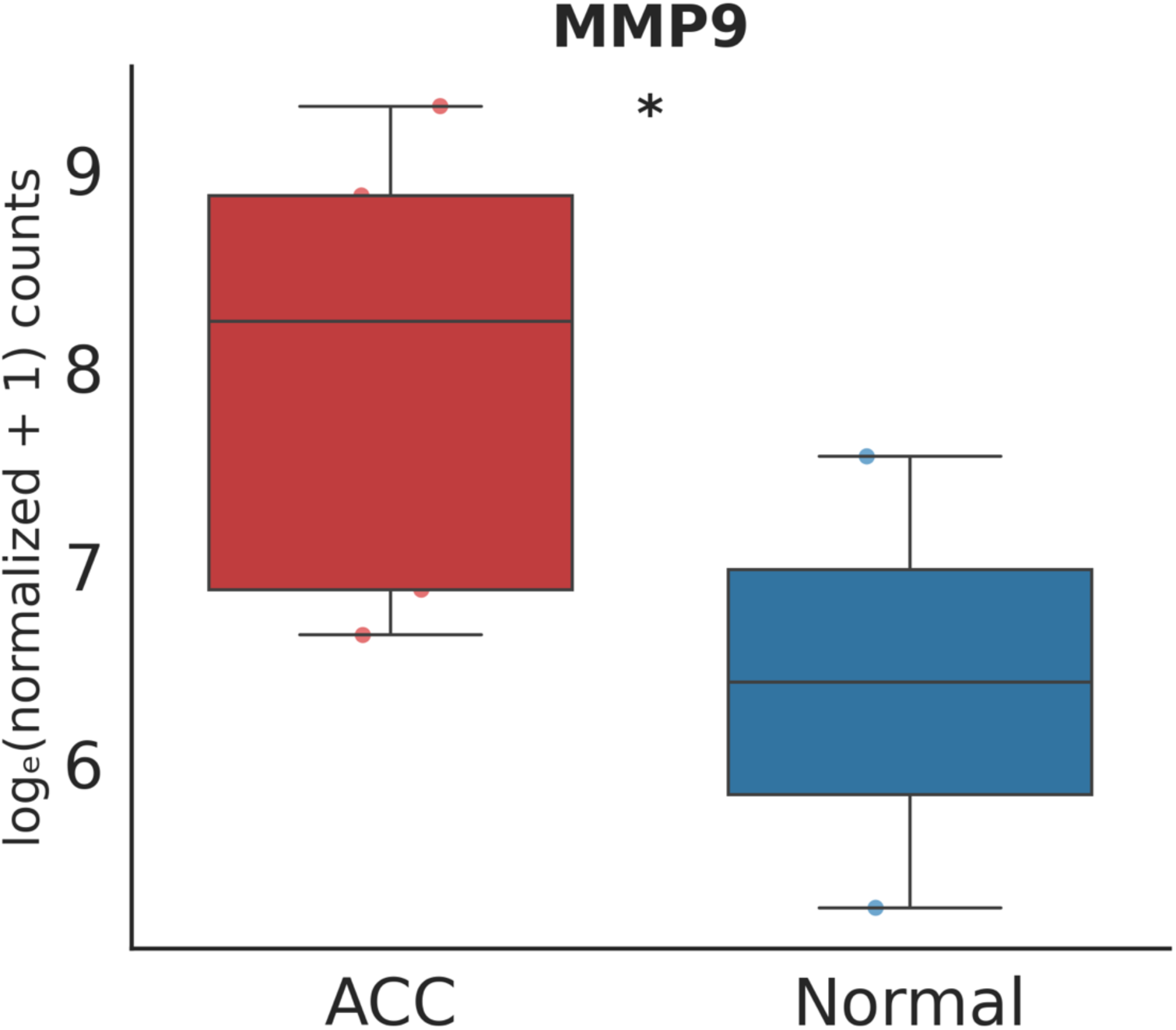
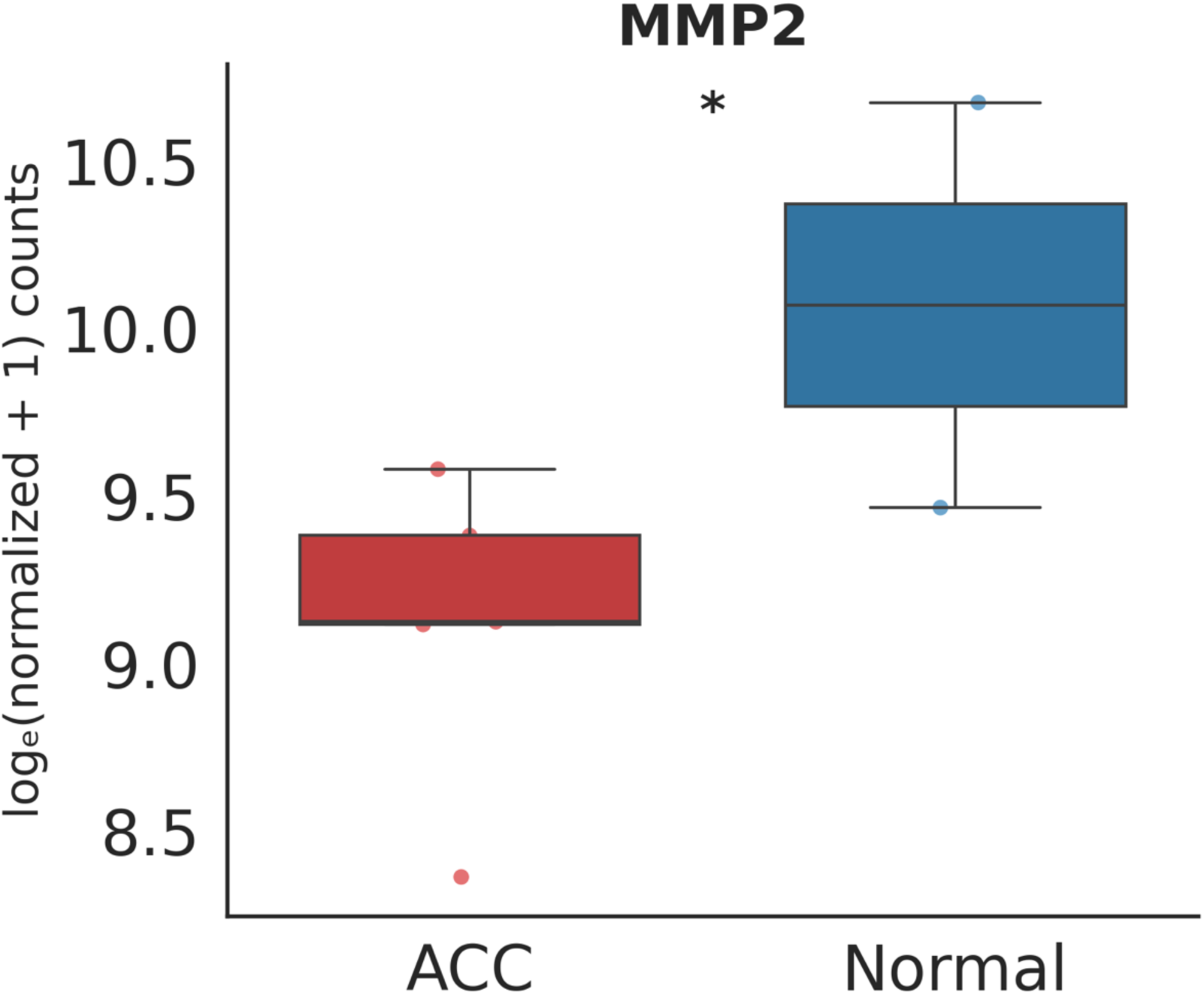
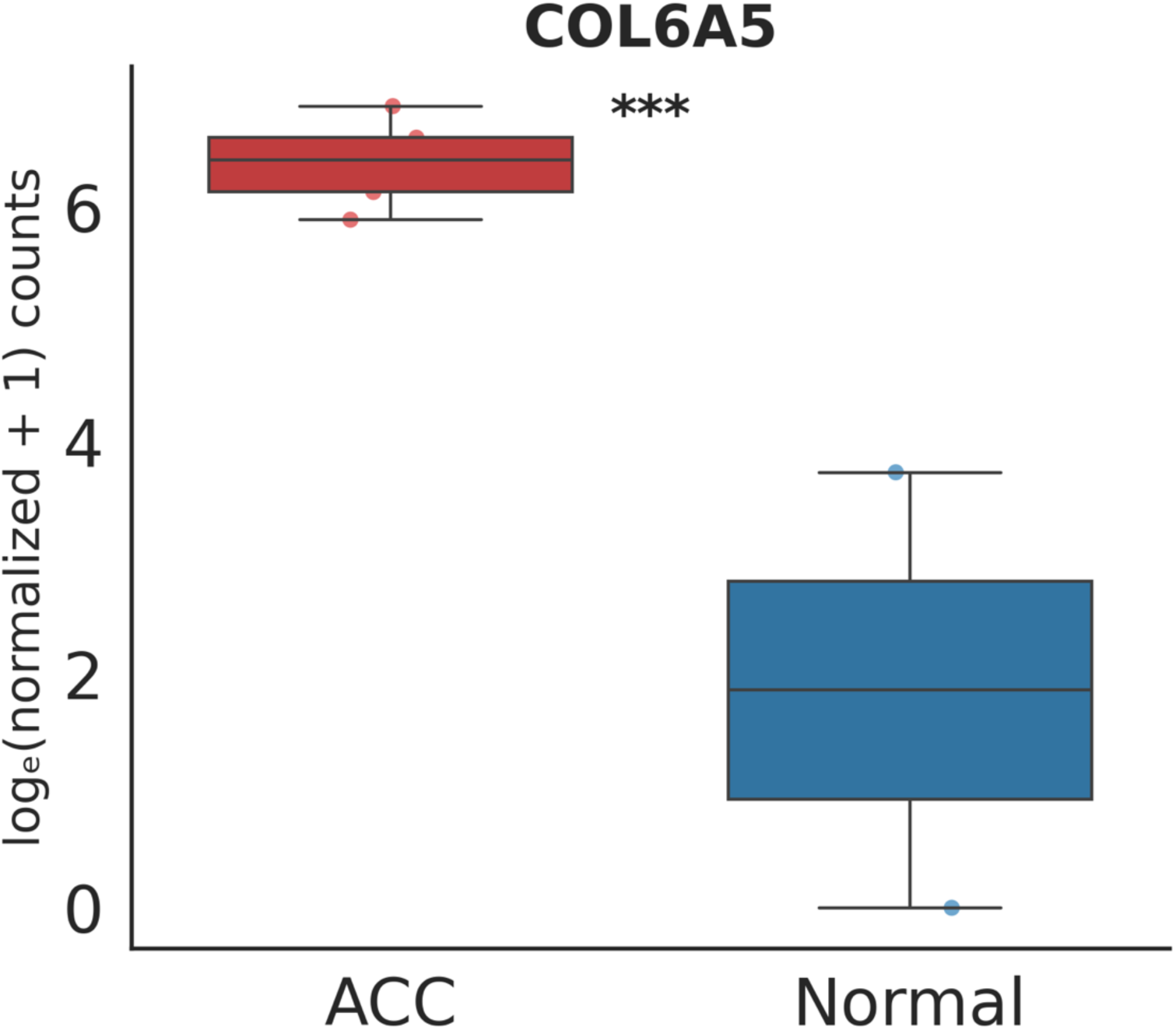
RNA sequencing analysis of spatial SGACC versus normal control salivary glands. (A) Heatmap of top 1000 differentially expressed genes of normal control (N1) samples (n =2) vs SGACC (ACC1 -ACC5) samples (n = 5). (B) Heatmap of top 60 differentially expressed genes of normal control (N1 and N2) vs SGACC (ACC1 -ACC5) samples (n = 7). Green and red bars indicate gene upregulation and downregulation, respectively. (C) Heatmap of top markers for each cluster considering 50 bins from each cluster. (D) **Box plots of top differentially expressed collagens and matrix metalloproteinases between normal salivary gland and SGACC.** Statistical significance is indicated by asterisks: * *P* < 0.05, \*\***P** < 0.01, and \*\*\***P** < 0.001.

### 2.3 Spatial Transcriptomics Analysis and Preprocessing

A customized computational pipeline, based on the methodology described by Xun Ding et al. [24], was used to extract unique mapped reads and perform spatial binning of the transcriptomic data. Initially, the query spatial SGACC dataset comprised transcripts from 19,833 genes across an estimated ∼10.5 million 2µm features, yielding a mean transcript count of 5.81 and a maximum of 629 transcripts per 2µm feature, along with an average of 5.48 and a maximum of 483 unique genes per 2µm feature. Spatial resolution was set at a bin size of 5 to accommodate a set of 5, 2µm features on the x and y axis of the chip, resulting in a capture area of 10 µm × 10 µm. Quality control (QC) variables were generated, and binned features with fewer than 1,000 transcripts were excluded, resulting in an updated QC distribution with a mean of 2,212 and a maximum of 10,979 transcripts per binned features, alongside a mean of 1,483 and a maximum of 5,441 unique genes per binned features (Fig. S1A - S1B). Subsequently, we filtered out genes that were expressed in less than 200 binned features while simultaneously filtering out binned features where less than 200 genes were expressed. Dimensionality reduction was performed using principal component analysis (PCA), retaining 25 principal components. Top 3000 highly variable genes were identified by using Scanpy’s (version = 1.10.3) HVG function^25^.

These top 3000 genes were further subsetted for Leiden clustering with a resolution parameter set to 0.3 yielding distinct cellular subpopulations which were further categorized as MYB-expressing and non-MYB expressing clusters as visualized in Uniform Manifold Approximation and Projection (UMAP) (Fig. 1A, 1B). The individual clusters of the cell populations were stratified by utilizing the Scanpy’s rank_genes_groups function to determine the top marker genes of each cluster. These clusters were annotated using several reference databases: the Human Cell Atlas, DAVID v6.8, and the Annotation of Cell Types (ACT)^21,26,27^. Additionally, trajectory inference was performed using Monocle3 (version 1.3.3) to investigate lineage relationships and differentiation pathways^28^.

### 2.4 Pathology Annotations

The H&E image of the spatial SGACC sample was annotated by a Board Certified Anatomical Pathologist (Fig. S5). Pathological annotations demonstrated a strong concordance between the identified cell types within the spatial tissue architecture and the UMAP-derived clusters from the transcriptomic analysis, confirmed through both manual review, computational clustering and cell2location model prediction.

### 2.5 Cell2Location

All spatial transcriptomics visualizations were generated using Cell2Location (version 0.1.3). Specifically, Cell2Location^29^ was employed to quantify gene expression profiles while preserving spatial localization, and enabling the construction of cell type distributions, spatial domain maps, and tissue architectural clusters (Fig. 3A–3C). Highly variable genes (HVGs) identified from integrated single-cell reference datasets were used to train the Bayesian Cell2Location model for spatial deconvolution, allowing high-resolution mapping of cell populations across the tissue. Cell2Location leverages reference-informed deconvolution to interpret gene expression patterns within spatial transcriptomics datasets, integrating prior knowledge from single-cell data (Fig. 1C), to enhance tissue-specific resolution. The robustness of the model was evaluated using the Evidence Lower Bound (ELBO) loss plot for quality control, and model validity.

**Figure 3.**
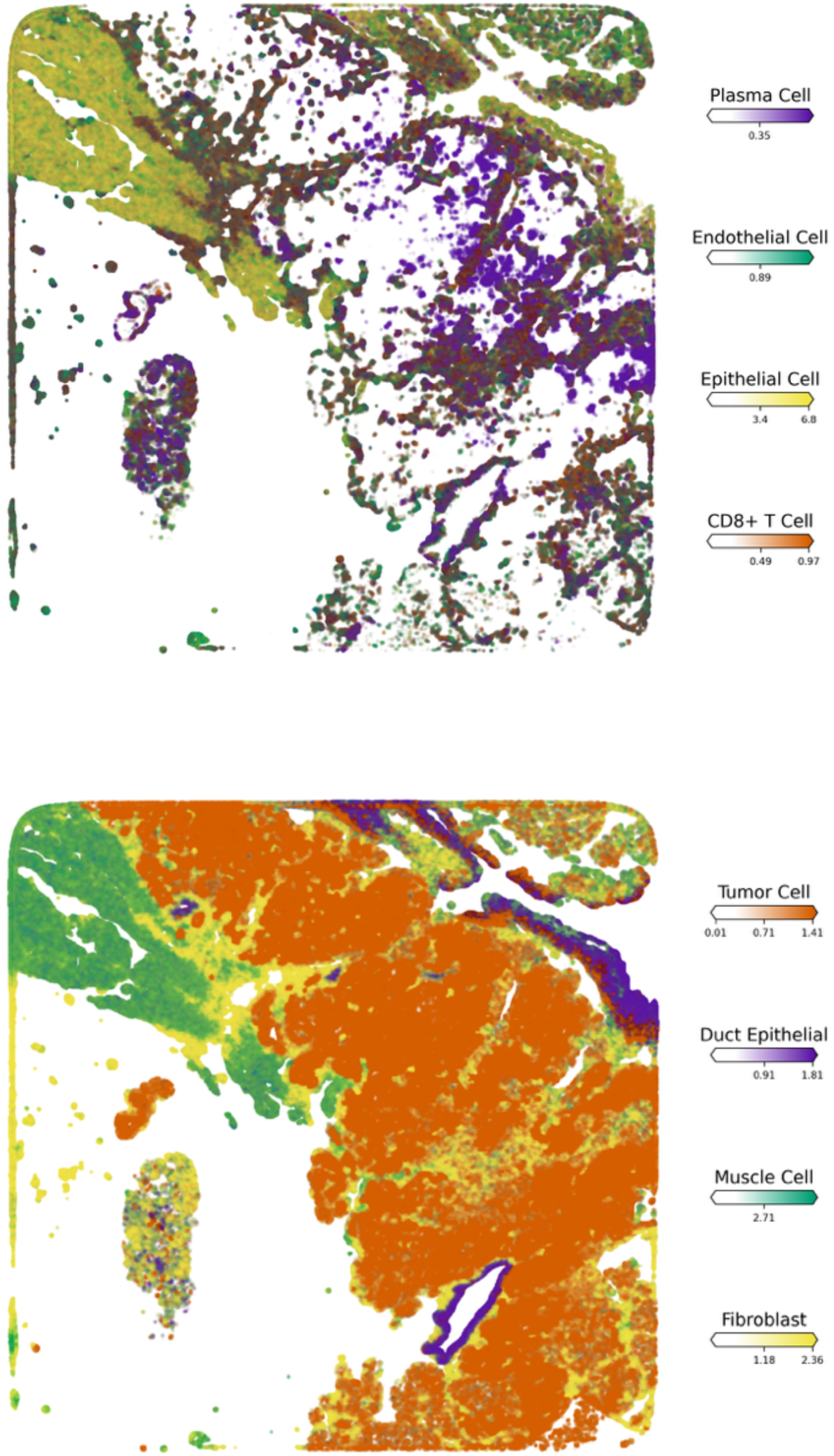
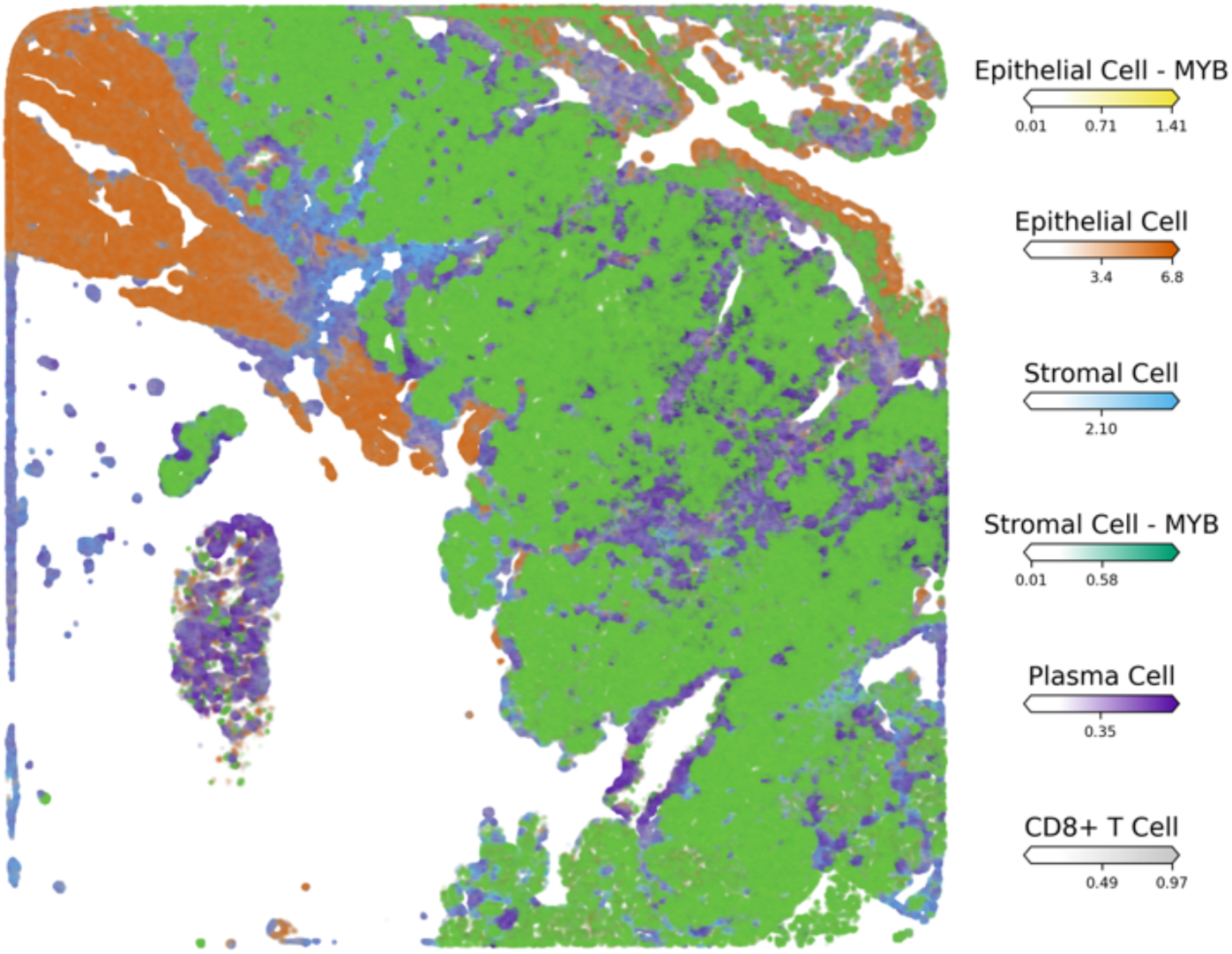
Identification of cell clusters in the spatial transcriptomics sample along with their relative abundance. (A). Spatial map showing absence of MYB-expressing cell clusters. (B). Co-localization of MYB-expressing cell clusters.(C). Visualization of intercellular crosstalk highlighting stromal cell populations

### 2.6 Gene Ontology (GO) and Pathway Analysis

Transcriptional regulation within MYB and non-MYB clusters was examined through gene ontology (GO) and pathway enrichment analyses. Differentially expressed genes were subjected to downstream analysis using the Database for Annotation, Visualization, and Integrated Discovery (DAVID) v6.8, employing GOTERM_DIRECT ontologies and KEGG pathway annotations. Visualization of GO terms and enriched pathways was performed using the *ggplot2* package in R^30^.

## 3.0 Results

### 3.1. Spatial Transcriptomics and Cell Cluster Annotation

To analyze the spatial architecture and cellular crosstalk between MYB and non-MYB tumoral cores, spatial transcriptomic profiling was performed on a Stage III–IV SGACC tissue sample from a patient with a recorded survival duration of 204 months. Histologically, this sample displayed a tubulocribriform architecture, a subtype recognized for its tendency to shift across histological grades I, II, and III (Table 1)^31^. The two-dimensional UMAP revealed 11 distinct transcriptional clusters, which were further stratified into four clusters demonstrated markedly elevated MYB expression, alongside with other transcriptionally diverse populations (Fig. 1A and B).

**Table 1:** Sample clinical Information.

| Clinical_information |  |  |  |  |  |  |  |
| --- | --- | --- | --- | --- | --- | --- | --- |
| Path | Copenhagen deidentifier | Translocation | STATUS | Survival Months | GENDE R | Form | Stage |
| 18-Rig70018 04.42.04 FORS UF SA42 | 18Rig750018044206 | MYB12-NFIB | Deceased | 125 | Female | Tubulocribriform | III-IV |
| 18-Rig750018; 04.39.04; FORS; UF; SA39 | 18Rig750018043906 | MYB14-NFIB | Alive | 175 | Male | Tubulocribriform | III-IV |
| 18-Rig750018; 04.39.05 FORS; UF; SA39 | 18Rig750018043906 | MYB14-NFIB | Alive | 175 | Male | Tubulocribriform | III-IV |
| 18-Rig750018; 04.42.05FORS; SA42 | 18Rig750018044206 | MYB12-NFIB | Deceased | 125 | Female | Tubulocribriform | III-IV |
| 18-Rig750018; 04.45.05 FORS; SA45 | 18Rig750018044506 | MYB8-Other | Alive | 204 | Male | Tubulocribriform | III-IV |

The heatmap of top marker genes (2C) was constructed by considering 50 bins from each cluster. The epithelial MYB-expressing cluster was characterized by elevated expression of canonical epithelial markers such as *AZGP1* and *ITGB4*. In contrast, the basal MYB-expressing cluster was delineated by the high expression of basal-specific markers including *KRT5* and *KRT17*. Endothelial cell populations were annotated based on the expression of *MGP* and *EGR3*. Non-MYB-expressing epithelial clusters were identified by their ubiquitous expression of markers such as *KRT13*, *SPRR2A*, *SBSN*, and *LY6D*. Fibroblast clusters were annotated by notable expression of extracellular matrix-related genes including *COL1A1*, *COL3A1*, and *COL1A2*. The muscle cell population was defined based on the corroborated expression of contractility-associated genes such as *TNNT3*, *CKM*, *MB*, *TTN*, and *STAC3*. Notably, all MYB expressing clusters were categorized as MYB expressing based on the expression of MYB.

Plasma cell (PC) clusters exhibited strong transcriptional signatures consistent with antibody-producing cells, including *IGKC*, *IGHA2*, *IGLC2*, and *JCHAIN*. Non-MYB stromal cells were annotated based on the expression of *CLEC3B*, *COL1A1*, and *HBA2*, whereas MYB-expressing stromal cells demonstrated high levels of *ALDH1A3*, *EGR3*, and *NR4A1*. These annotations were further validated through complementary molecular and biological relevance assessments, and pathway enrichment analyses (Table S1).

### 3.2. Zonal Distribution of MYB Expressing Clusters Exhibit Spatial Co-localization within Tissue Architecture

Following the classification of cell clusters based on MYB expression levels within the cellular population, we examined the spatial distribution of MYB-expressing versus non-MYB-expressing cell states and the spatial mapping revealed a distinct co-localization of all MYB-expressing clusters including Basal-MYB, Stromal-MYB, and Epithelial-MYB subtypes which are predominantly structural in nature. These clusters exhibited tight spatial association, in contrast to non-MYB-expressing clusters that demonstrated a zonal distribution that is localized more distally from MYB-expressing populations which largely occupied the interior regions of the tissue architecture (Fig. 3A-C). This pattern suggests underlying biological and pathological commonalities among MYB-expressing clusters.

The observed co-localization of MYB-positive cells may reflect the presence of a dominant oncogenic clone driven by MYB-NFIB gene fusions, which are well-established molecular hallmarks of tumorigenesis in SGACC (Fig. S2A-B & S3A -B). Such clonal expansions likely result in compartmentalization, wherein transcriptionally similar cancer cells proliferate and aggregate within defined tumor niches, consistent with a shared oncogenic program and lineage derivation from a transformed progenitor population^8,32,33^. Additional factors potentially contributing to this spatial organization include extracellular matrix remodeling, altered cell-cell adhesion dynamics, partial activation of epithelial-mesenchymal transition (EMT) programing, recruitment of reactive stromal elements, angiogenesis, and modulation of local immune cell populations which are observed to be significantly up-regulated in the GO analysis. These processes may cooperatively shape the tumor microenvironment (TME), facilitating the establishment of pro-tumorigenic niches^34,35^.

Furthermore, the spatial behavior of MYB-expressing populations may be influenced by structural constraints conferred by the presence of ducts and stromal compartments. These physical and molecular feature together with signaling pathways such as PI3KCA and an aberrant dysregulated Wnt signaling likely contribute to the infiltrative and invasive characteristics typical of SGACC tumors^36^. All these observations are corroborated by the pathology annotations and significantly upregulated biological processes and pathway analyses (Fig 4A-C). This colocalization is evidently confirmed by the empty center-region of the spatial visualization (Fig 3A). The spatial localization of individual cells is presented in the supplementary materials (Fig. S7)

**Figure 4.**
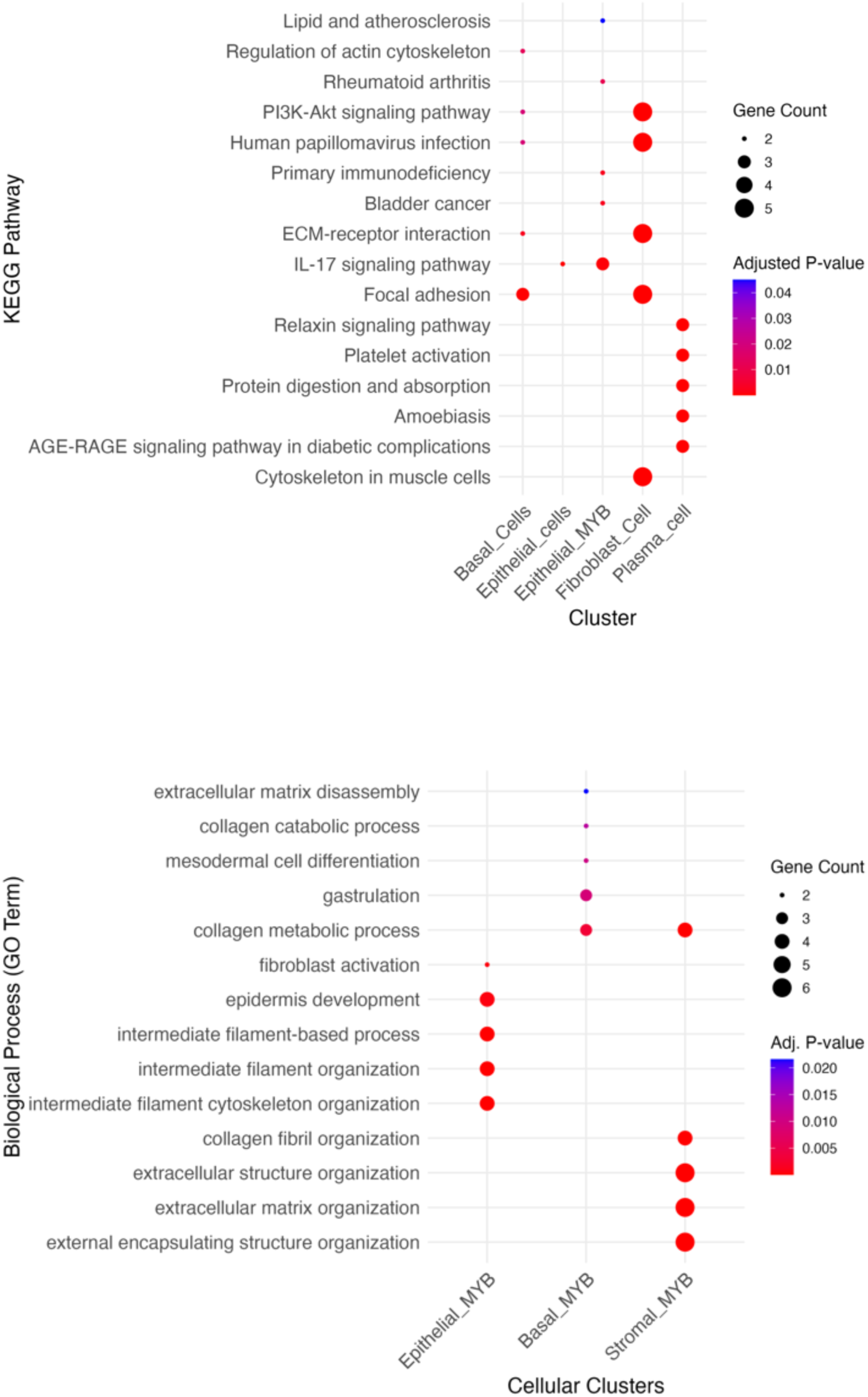
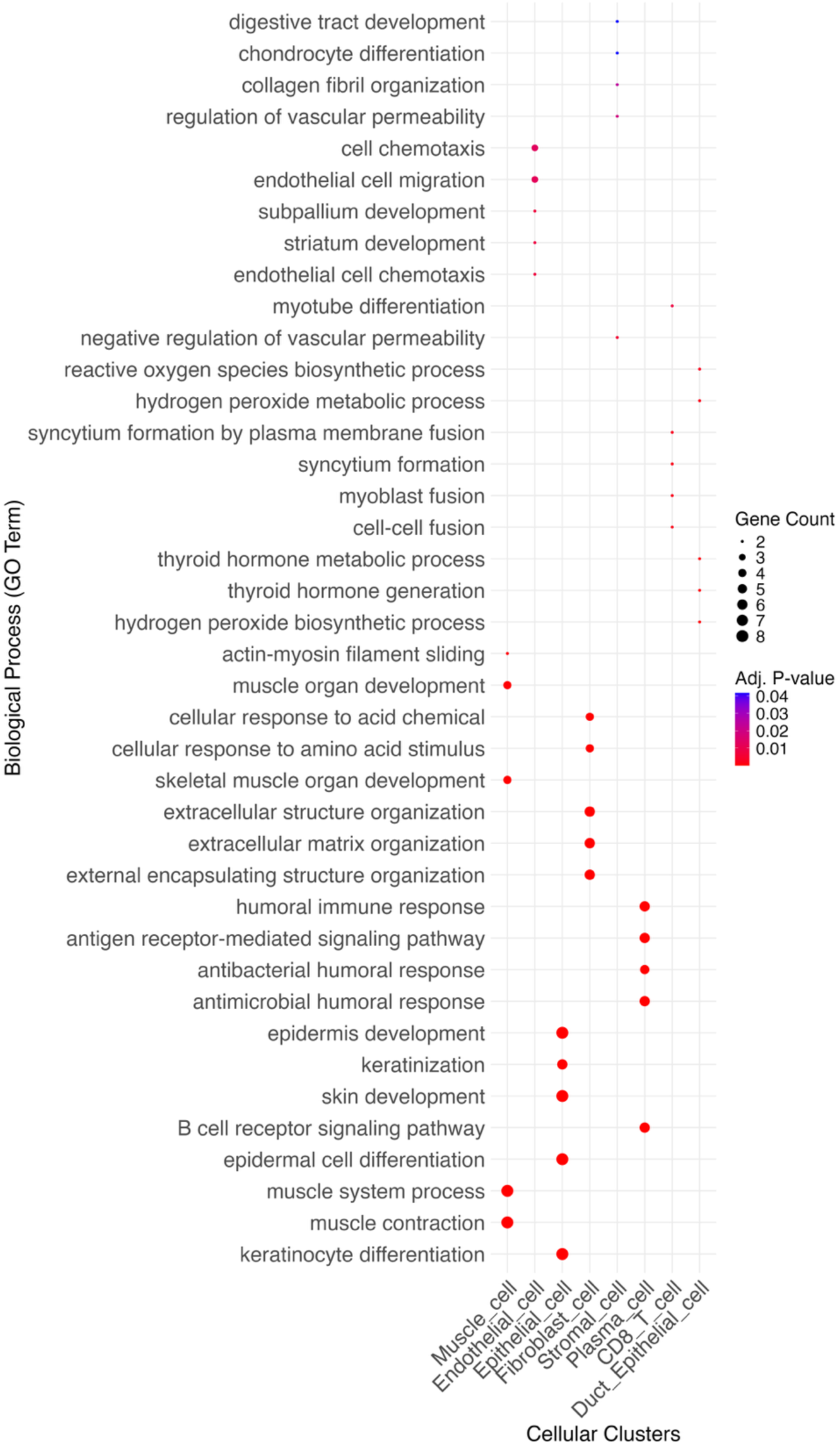
Kegg pathway and biological process. (A) KEGG pathway enrichment analysis in MYB-expressing and non-MYB-expressing clusters showing IL-7 and PI3K pathway upregulation (B). GO biological process enrichment analysis in MYB-expressing clusters. (C). GO biological process enrichment in non-MYB-expressing clusters.

### 3.3. Spatial Visualization highlights a localized enrichment of immune cells within the SGACC tissue microenvironment

To study the distribution of immune cells within the tumor microenvironment, we employed differential gene expression analysis alongside spatial cellular deconvolution techniques. Our findings show that immune cell infiltration into tumor niches predominantly occurs through regions enriched with MYB-expressing structural cells (Fig. 5 A-C). Spatial transcriptomic analysis revealed a localization of CD8⁺ T cells within the MYB and non-MYB expressing cell populations (Fig. 5A), further demonstrating a consistent colocalization of the immune cell cluster with the tumoral core. This infiltration within the tumor is evidenced by both quantitative abundance (Fig. 5D) and spatial distribution which is widely recognized as a positive prognostic marker and reflects a tumor immune microenvironment (TIME) conducive to effective immune checkpoint inhibition (ICI). This immune infiltration may in part account for the prolonged patient survival exceeding 240 months, as recorded in the clinical metada.^37,38^

**Figure 5.**
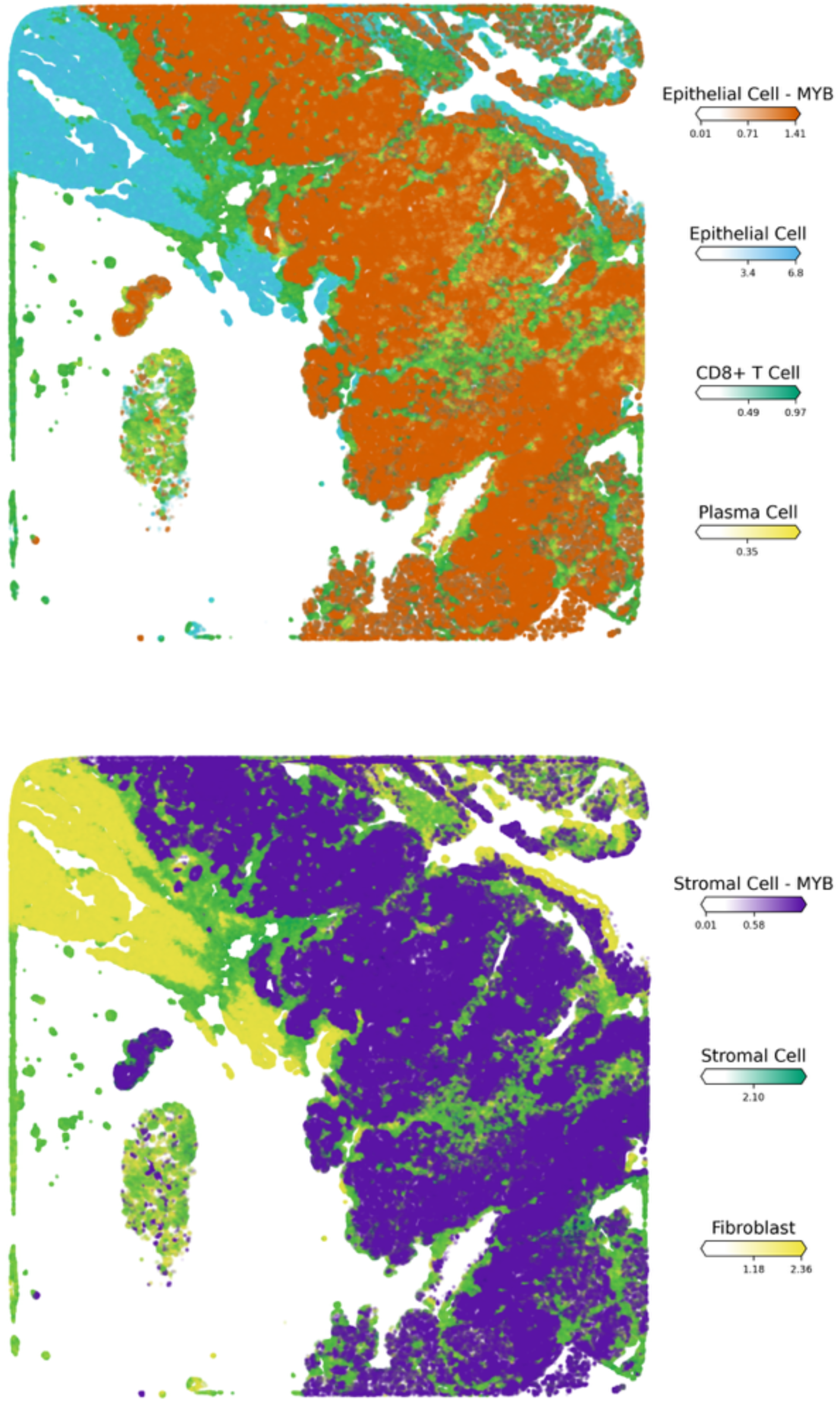
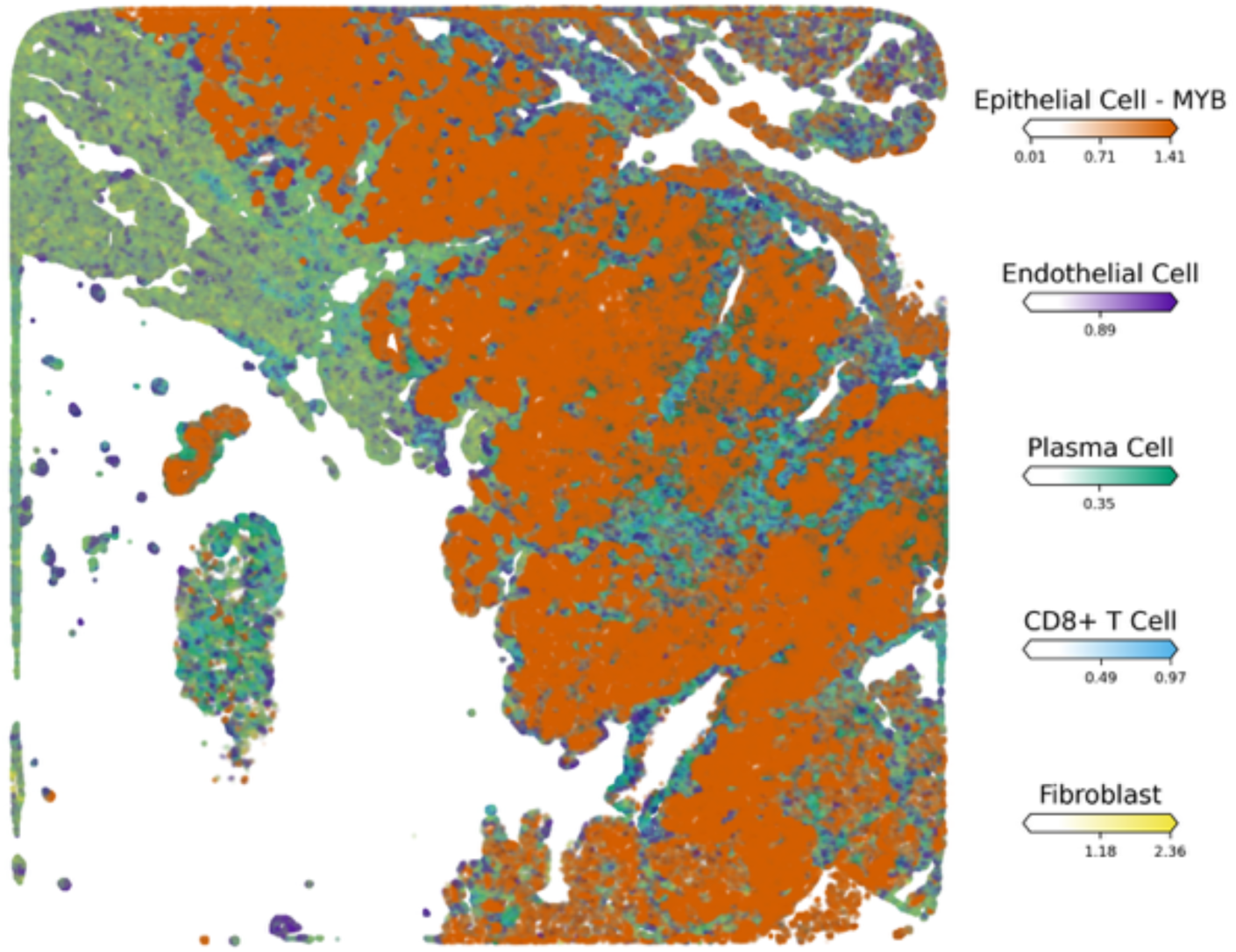
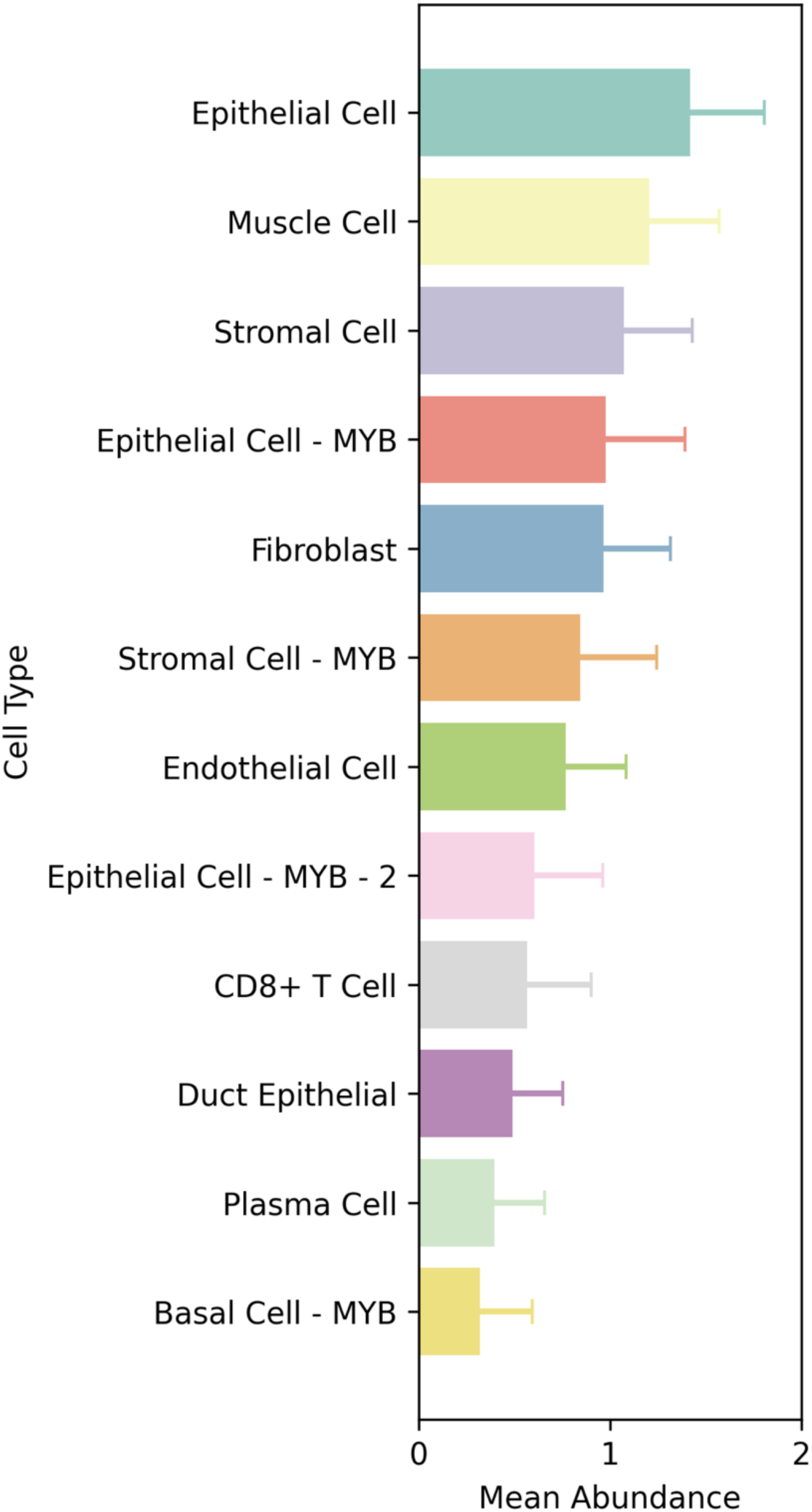
Identification of immune cells interactions with cell populations in the spatial transcriptomics sample along with their relative abundance. (A) Visualization of plasma and T cell interactions with MYB-expressing clusters. (B) Spatial distribution of an active stromal cell population within the tumor core. (C) Plasma cell migration toward tumor regions along fibroblastic and endothelial scaffolds (D) Spatial abundance of annotated cell types across the tissue section

**Figure 6.**
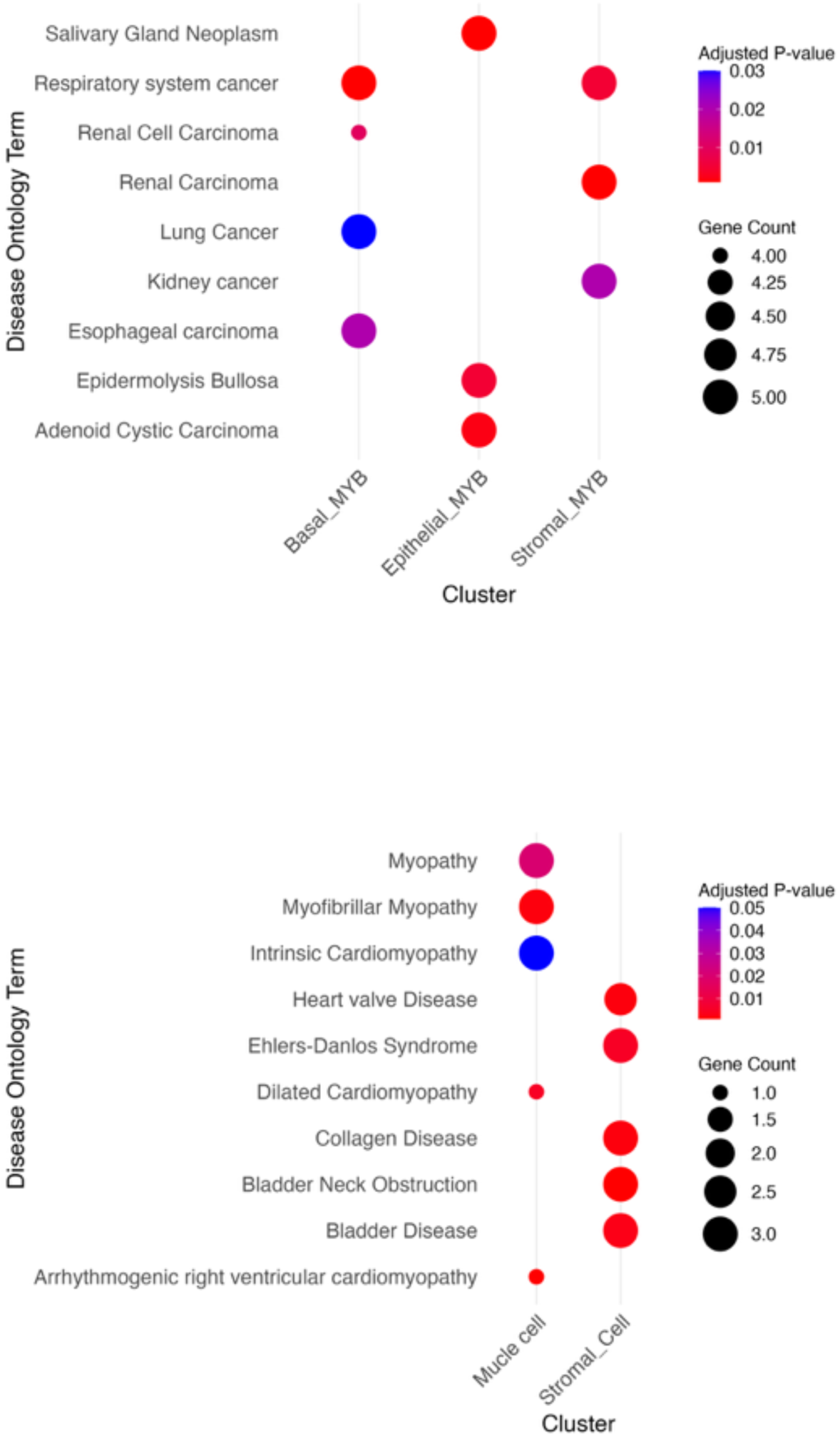
Enrichment of disease-related pathways and angiogenic processes. (A). Upregulated disease-associated pathways in MYB-expressing cell clusters. (B) Upregulated disease-associated pathways in non-MYB-expressing cell clusters.

The Gene Ontology (GO) analysis of the T cell cluster revealed a marked upregulation of pathways associated with αβ T cell activation and cellular components related to the αβ T cell receptor complex, indicating a heightened antigen recognition and active engagement with antigen-presenting cells (APCs)^37,39,40^. Interestingly, the molecular function category also showed a significant enrichment of heparin binding, suggesting enhanced T cell adhesion properties and extracellular matrix remodeling features consistent with adaptation to a dynamically altered TME. This may further reflect a state of T cell exhaustion, potentially resulting from sustained antigenic stimulation over time^41–48^.

Critically, spatial transcriptomic analyses demonstrated that plasma cells are colocalized with MYB-expressing tumor cell populations, indicative of an endogenous anti-tumor immune response. We observed expression of tertiary lymphoid structure (TLS)-associated gene signatures including *CD79A*, *CXCL8*, and *IL7R* (interleukin-7 receptor) which supports the presence of well-organized TLS within the tumor microenvironment. The TLS appears to facilitate directed plasma cell migration toward tumor regions along fibroblastic and endothelial scaffolds (Fig. 5C) ^37,49^.

Furthermore, this spatial and transcriptional immune architecture was accompanied by significant upregulation of the B cell receptor (BCR) signaling pathway and a concomitant increase in the expression of immunoglobulin-related components. These include the IgM-BCR complex, secretory IgA complex, IgD and IgG complexes, and general immunoglobulin-associated structures. The enrichment of antigen binding and immunoglobulin receptor binding functions suggests an active humoral immune response, possibly indicative of a *de novo* or sustained response to ICI by the patient’s immune system^8,50^. It is worth noting that spatial differential analysis showed that genes for plasma cells and CD8+ T cells were not differentially expressed between MYB- and non-MYB-expressing regions. This indicates a relatively comparable pattern of immune cell distribution across areas of the tissue architecture (Supplementary Documents 4 and 5).

### 3.4. Cellular Interaction Analysis Reveals Upregulated Disease Pathways, Angiogenesis, and Vascularization in SGACC

To interrogate biologically significant pathways associated with spatial interactions between MYB- and non-MYB-expressing cell clusters, functional enrichment analyses were conducted using gene ontology enrichment analysis along with spatial conformation from Cell2Location analyses. Clusters which were considered for pathway upregulation were examined to identify disease-relevant signaling pathways beyond the traditionally cited canonical mechanisms.

Interestingly, clusters with elevated MYB expression exhibited significant enrichment for disease-associated pathways such as bladder cancer and primary immunodeficiency. In addition to these, pathways specific to ACC and MYB neoplastic proliferations such as, malignant neoplasms of the salivary glands, and colorectal carcinoma were also detected. Gene Ontology (GO) enrichment further revealed robust upregulation of angiogenesis-related transcriptional programs. This was exemplified by the elevated expression of vascular and endothelial markers including *VEGFR2*, *VEGFC*, *XPNPEP2*, *THBS4*, *LEPR*, and *VCAM1*, MMRN1 suggesting an active vascular remodeling^51–53^.

These transcriptional changes are likely driven by the metabolic and oxygen demands of proliferating tumor cells in a hypoxic and an expected immunosuppressive microenvironment. In such contexts, neovascularization becomes essential to support tumor progression by enabling enhanced delivery of nutrients and oxygen. Furthermore, the observed activation of angiogenic pathways is consistent with the angiogenic switch, a critical step in tumor progression that marks the transition from an avascular to a vascularized state. This switch is frequently coordinated with EMT, wherein hypoxia-induced signaling promotes EMT in tumor-associated endothelial cells, thereby facilitating tumor invasion, endothelial chemotaxis, and metastasis^54,55^.

### 3.5. Monocle Trajectory and Pseudotime Analysis reveals partial EMT programming

We conducted trajectory inference and pseudotime analysis using the Monocle3 framework, focusing on cells exhibiting significant upregulation of the *MYB* gene. The analysis encompassed the highest *MYB*-expressing clusters originating from epithelial, basal, and stromal lineages, with the objective of identifying a potential tumor-initiating population and tracing the lineage relationship among these cell types. This approach aimed to identify the cluster most analogous to the progenitor and Induced Pluripotent Stem Cells (IPSCs), potentially facilitating the proliferative cancer cell expansion across the tissue (Fig. 7 A-D).

**Figure 7.**
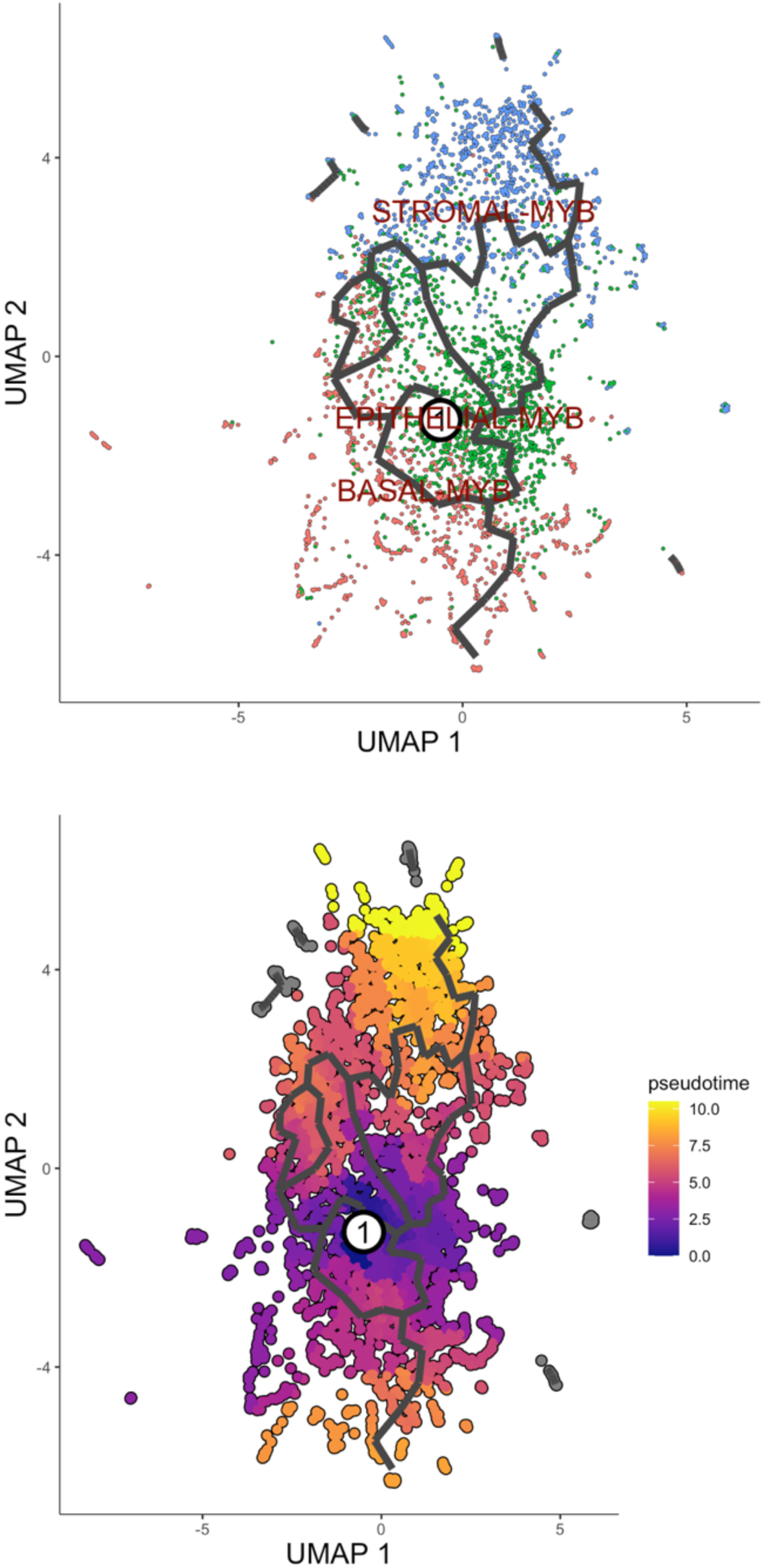
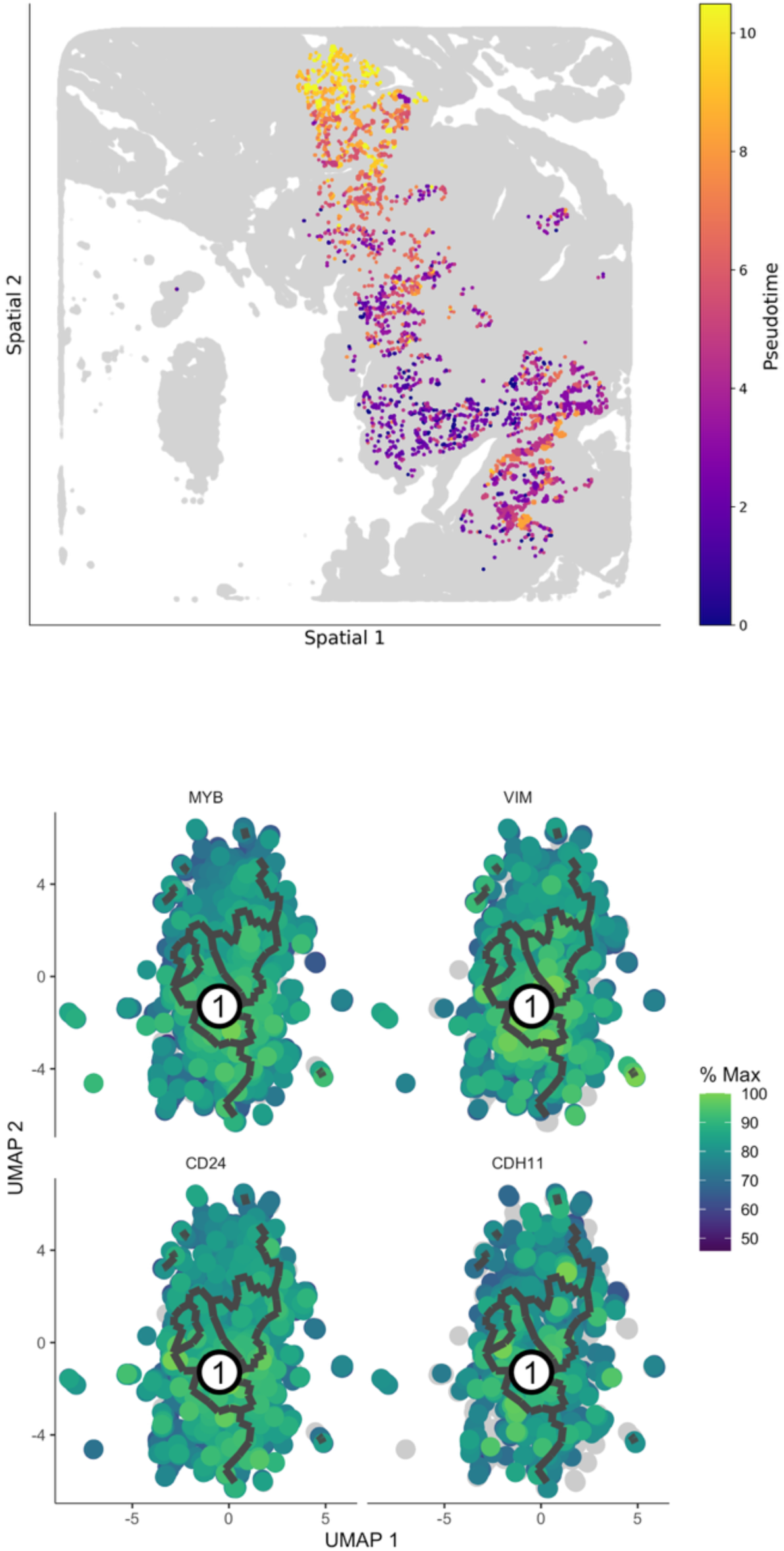
Trajectory and pseudotime analysis of MYB-expressing cell clusters. (A). Monocle-inferred trajectory of key MYB-expressing cell clusters. (B). Pseudotime estimation across principal MYB-expressing populations. (C). Spatial mapping of pseudotime progression within the tissue architecture. (D) UMAP visualization of selected genes identified from trajectory analysis.

The pseudotime mapping revealed that the epithelial *MYB*-expressing cluster constitutes the root cell population, from which the stromal and basal *MYB*-expressing cells appear to derive. Independent enrichment analyses were performed on each of these clusters to identify distinct pathway enrichments (Fig. S4-GO of the 2 groups of MYB clusters). Notably, the epithelial *MYB* cluster demonstrated marked upregulation of gene programs associated with intermediate filament organization, epithelial cell differentiation and a pronounced expression of vimentin which is the hallmarks of epithelial-mesenchymal transition (EMT). However, comparative analysis with non-*MYB*-expressing epithelial populations revealed the retention of epithelial features in the latter, including robust cell-cell adhesion and keratinocyte differentiation. These phenotypes were supported by elevated expression of canonical epithelial markers such as *CRNN*, *CSTA*, *ANXA9*, and *DSG3*. We further present a visualization of gene expression dynamics for *MYB*, *CD24*, *CDH11*, and *VIM* along with the spatial pseudotime trajectory (Fig. 7D), highlighting the progressive transcriptional shifts asso1ciated with tumor progression and EMT^56^. During the EMT programing, epithelial cells adopt mesenchymal characteristics which results in a transitional state that facilitates the directional movement of cancer stem cells (CSCs), that are implicated in tumor initiation, metastasis, therapeutic resistance, and disease recurrence. Experimental studies have demonstrated that subpopulations of CSCs derived from mammary epithelial cultures can form mammospheres upon stimulation with TGF-β, underscoring their self-renewal capacity and role in cancer progression^57^.

However, if EMT programming were not observed from the trajectory inference, the epithelial cells would have exhibited a characteristic apical-basal polarity and maintain close contact with neighboring cells through specialized junctional complexes, including adherens junctions, tight junctions, and desmosomes. In contrast to mesenchymal cells which are typically embedded within the extracellular matrix and lack defined apical-basolateral polarity^55,58^.

Analysis of the three most prominent MYB-expressing clusters revealed phenotypic features indicative of mesenchymal traits, supporting a transition consistent with partial epithelial mesenchymal transition (P-EMT). This hybrid epithelial-mesenchymal state is frequently associated with the acquisition of aggressive carcinoma phenotypes. Our trajectory analysis revealed evidence of extensive trans-differentiation among the cell populations, marked by pronounced differentiation from progenitor to mature mesenchymal states, loss of epithelial cell-cell adhesion, and partial retention of focal adhesion features, particularly in basal MYB-expressing cells. Additionally, reactive stromal cells exhibited signs of keratinization, while some cells showed a reduction in keratin expression which are hallmarks consistent with P-EMT. The presence of aberrant negative regulation of canonical Wnt signaling in a lower MYB expressing cells adjacent to the stromal compartment and some immune cells further supports the occurrence of P-EMT^56^.

Our findings indicate a mesenchymal progression originating from clonal epithelial cells, culminating in a differentiated mesenchymal subpopulation with the highest pseudotime value of 10, while the root epithelial progenitors exhibited a pseudotime of zero. This trajectory suggests a developmental lineage from epithelial to mesenchymal states and are consistent with previous reports in invasive ductal carcinomas, in which transformed stromal cells harbor chromosomal rearrangements shared with epithelial tumor cells, implying a common clonal origin and the retention of stroma-specific genomic alterations even at advanced differentiation stages, including the IPSC-like state^59–61^.

Although we did not observe pronounced expression of canonical EMT-inducing transcriptional drivers such as TWIST, we uncovered the presence of other transcriptional drivers such as ZEB1, SNAI1, SNAI2. Also, the expression of the mesenchymal marker Vimentin, along with multiple microenvironmental cues within the SGACC tumor, suggests the initiation of a mesenchymal transition program (MTP). These cues include key EMT-related signaling pathways mediated by transcriptional regulators such as TGF-β, HIF-1α, EGF, WNTs, and Notch. This pattern points to a context-dependent and selective activation of EMT-associated transcription factors, likely driven by heterotypic interactions with neighboring cancer cells and tumor-associated stromal elements. Such interactions may facilitate the emergence and expansion of cells with specialized mesenchymal phenotypes, contributing to local invasion and phenotypic diversification within the tumor microenvironment^58^.

### 3.6. Pairwise analysis between MYB and NON-MYB clusters - Differential Expression

In our differential gene expression analysis aimed at understanding molecular distinctions among the three principal MYB-expressing clusters in the spatial SGACC, we compared these with their respective non-MYB-expressing counterparts across stromal, epithelial, and basal cell populations. These specific cell populations were prioritized based on insights from pseudotime trajectory analysis, which identified MYB-expressing epithelial cells as potential tumor-initiating clusters. Moreover, stromal cells were characterized as the terminal state of the P-EMT programing, reflecting their acquisition of functional attributes associated with mesenchymal differentiation and the subsequent tumor proliferation driven by the presence of diverse Cancer Stem Cells (CSCs). To quantify differential expression, we statistically tested 3,000 genes for each pairwise comparison between MYB-expressing and non-MYB-expressing populations using two sample t-tests. Genes with a *P*-value < 0.05 were considered significant. To correct for multiple hypothesis testing, we applied the Benjamini–Hochberg FDR to control the false discovery rate (FDR).

Upon concluding the pairwise computation, we analyzed a total of 30 differentially expressed (top 15 upregulated and top 15 downregulated) genes in the MYB expressing cell types for each comparison. For the stromal cell comparison, our analysis revealed upregulation of genes associated with altered epithelial states and reactive stromal activity, exemplified by the increased expression of *Kruppel-like factor 6 (*KLF6) and CLCA2. We also observed elevated expression of transcriptional signatures implicated in key oncogenic signaling pathways, including those promoting tumor progression and lymphatic metastasis (Fig. 8A). Several genes associated with adaptive responses to cellular stress, focal adhesion remodeling, and enhanced tumor cell survival were also upregulated. Intriguingly, we noted ectopic expression of genes such as TCTE1, TRAF1 which are typically restricted to gonadal tissues, suggesting aberrant activation of survival mechanisms in ACC. Additionally, upregulation of genes involved in inflammation, polyamine catabolism (potentially contributing to reactive oxygen species generation), and simultaneous promotion of both cell adhesion and migratory behavior was observed. These changes, highlighted by the upregulation of KIF1B, VEGFC, BTG2, DOCK4, SHCBP1L, ATF3, and SPRR2E, underscore the tumor’s capacity for invasion, metastasis, and adaptation within a dynamic TME^62–69^.

**Figure 8.**
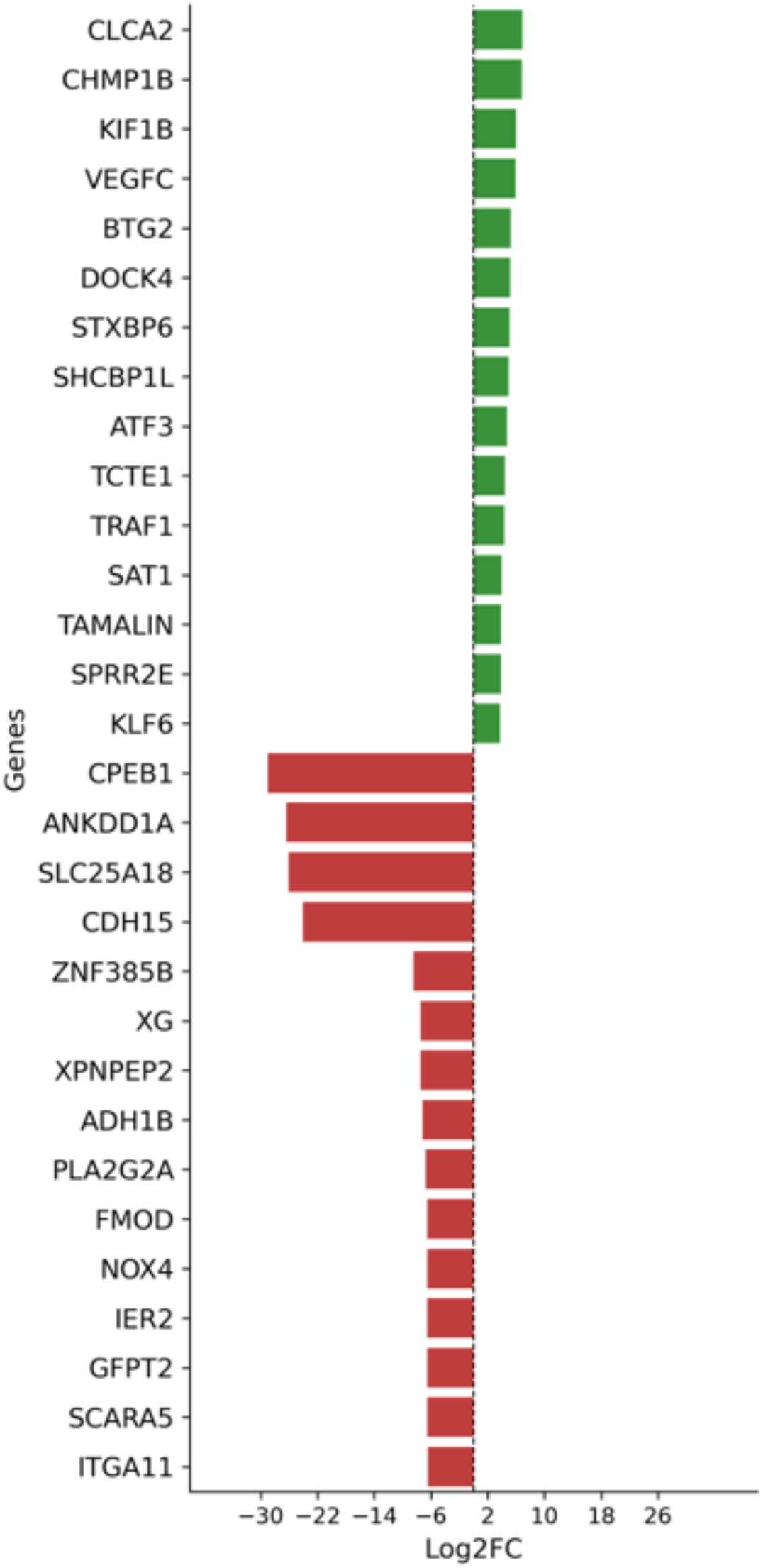
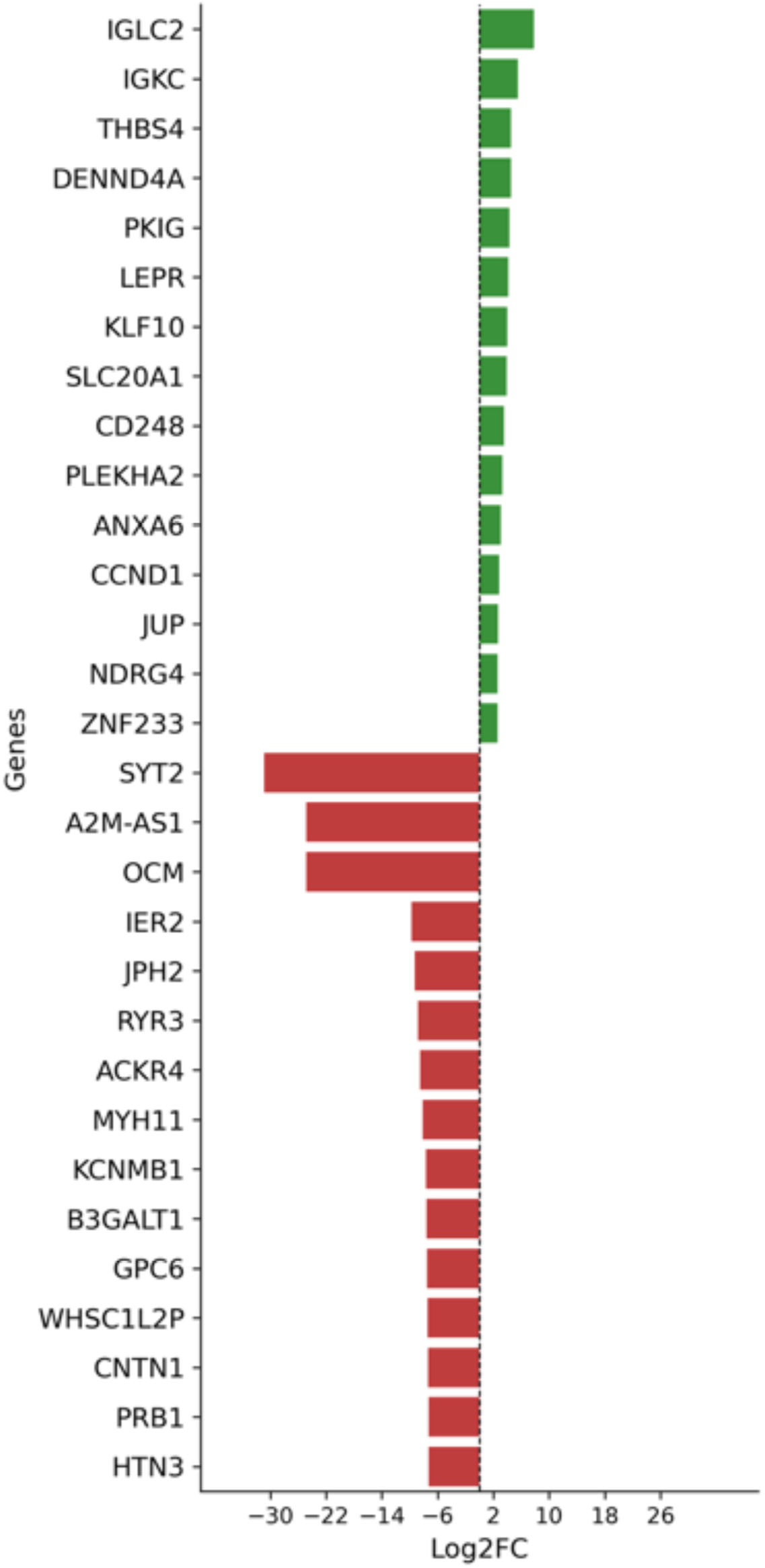
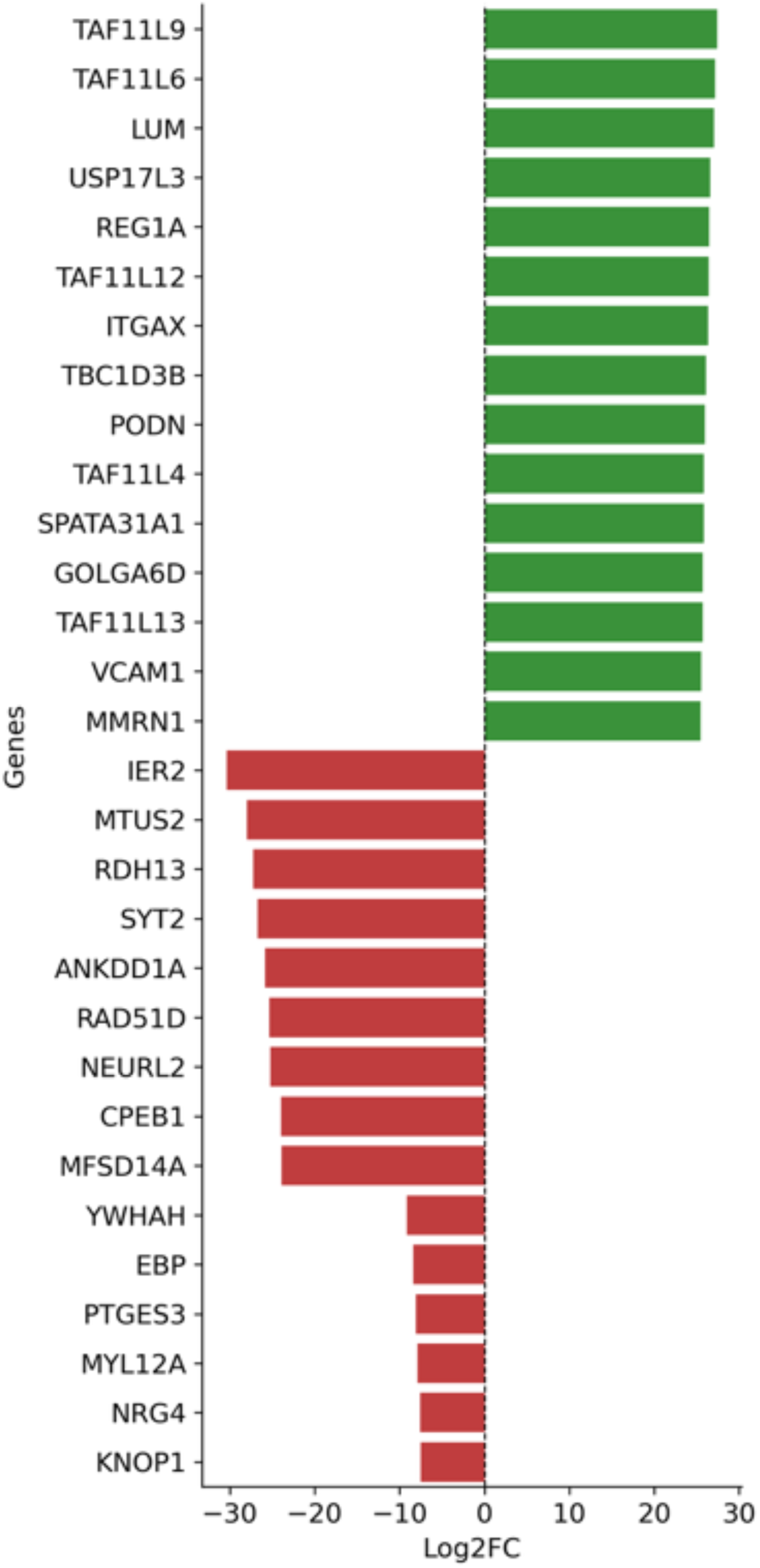
Differential gene expression between MYB and non-MYB expressing cell types. (A) Pairwise comparison of stromal cells expressing MYB vs. non-MYB. (B) Pairwise comparison of epithelial cells expressing MYB vs. non-MYB. (C) Pairwise comparison of basal cells expressing MYB vs. non-MYB.

Conversely, we observed a significant downregulation of tumor-suppressive transcriptional regulators such as *CPEB1*, CDH15, *GFPT2*, and *PLA2G2A*. This was accompanied by decreased expression of genes involved in cellular adhesion and differentiation, suggesting a loss of structural integrity and cellular identity. Notably, the reduced expression of *XPNPEP2* and *XG* implies heightened vascular permeability and an inflammatory microenvironment, which may further facilitate tumor progression. The downregulation of *FMOD* suggests a weakening of the stromal barrier, potentially enabling tumor invasion. In addition, diminished expression of *IER2* indicates an impaired stress response pathway, allowing ACC cells to circumvent critical cellular stress checkpoints. Finally, the decreased expression of *SCARA5* and *ITGA11* is indicative of enhanced cell detachment and invasive potential, further supporting a pro-metastatic phenotype^70–78^.

In the epithelial pairwise comparison, we observed a significant upregulation of IGLC2 and *IGKC*, suggesting an active immune response initiated at the root epithelial cell population (Fig. 8B), where tumor proliferation appears to trigger EMT programming. Furthermore, we identified upregulation of genes such as *THBS4*, *DENND4A*, *PKIG*, *LEPR*, CD248, and *PLEKHA2*, all of which have been implicated in key oncogenic processes, including angiogenesis, cell migration, invasion, extracellular matrix remodeling, activation of cancer associated fibroblast and tumor progression. Interestingly, we also observed increased expression of KLF10 and JUP, genes typically associated with tumor-suppressive activity and cell-cell adhesion, respectively. Additionally, *NDRG4*, a gene known for its role in mediating cellular stress responses, was also upregulated, suggesting a complex interplay between oncogenic signaling and stress adaptation within the tumor microenvironment^79–88^.

Conversely, the genes downregulated in the MYB-expressing epithelial clusters were predominantly tumor-suppressive transcriptional signatures, including *OCM*, *IER2*, *JPH2*, *RYR3*, MYH11and *B3GALT1*. These genes are broadly associated with rapid and transient cellular responses to growth factors and stress, regulation of controlled proliferation, maintenance of myoepithelial differentiation, inhibition of tumor cell migration, and preservation of cell-cell adhesion. Their reduced expression may contribute to the dysregulation of these critical pathways, thereby promoting tumor progression^89–94^.

In the basal pairwise comparison, we observed the upregulation of transcriptional signatures such as LUM, USP17L3, REG1A, TBC1D3B which are associated with remodeling of the extracellular matrix, facilitation of tumor invasion and metastasis, tumor survival, proliferation, enhancement of proliferative capacity and the alteration of signaling pathways and cell matrix interactions in tumor cells along with the upregulation of VCAM1 and MMRN1 which are fingerprinted to be associated with recruitment of cells involved in vascularization and promotion of extravasation of tumor cells by promoting their adhesion to tumor endothelial cells indication of pointed angiogenic process in the basal cells. Another paradoxical phenomenon is the upregulation of ITGAX which are associated with a possible influence on immune cell interactions within the tumor microenvironment, potentially affecting immune surveillance and tumor immunity^95–102^. All these along with upregulation of several pseudogenes (TAF11) and their variants. These pseudogenes may play unrecognized regulatory roles in gene transcription, warranting further investigation. Should these elements function as active paralogs or modulators, they may contribute to the transcriptional reprogramming of ACC cells, potentially promoting pro-proliferative, pro-survival, and pro-invasive phenotypes.

We observed a downregulation of several key genes, including NRG4, MYL12A, EBP, YWHAH, MFSD14A, CPEB1, NEURL2, RAD51D, ANKDD1A, SYT2, RDH13, IER2, and MTUS2. The reduced expression of these genes has been associated with various oncogenic processes such as resistance to apoptosis, sustained proliferation, increased oxidative stress, loss of tumor suppressor function, and dysregulated immune and inflammatory responses that collectively support tumor progression. Additionally, the downregulation of CPEB1 and related factors may lead to alterations in translational control that promote oncogenic protein synthesis, while the repression of MYL12A and IER2 is indicative of a loss of myoepithelial differentiation and cell identity, confirming the maturation of the EMT programming.

Interestingly, we also note the downregulation of PTGES3, a gene theoretically involved in supporting tumorigenesis, suggesting a paradoxical molecular signature that may reflect complex regulatory feedback, context-dependent tumor behavior or tissue heterogeneity^90,103–114^.

## 4.0 Discussion

Adenoid cystic carcinoma (ACC) of the salivary gland is distinguished by pronounced histological and molecular heterogeneity, rendering its underlying biology difficult to deconvolute and posing substantial challenges for its study in isolation. In this study, we employed single-cell RNA sequencing integrated with spatial transcriptomics to analyze the tumor architecture. Through a comparative analysis of the core MYB-expressing niche and adjacent non-MYB-expressing cell populations, we identified mechanistic drivers contributing to the observed tumor neoplastic cellular diversity. Our findings offer novel spatial and transcriptional insights into the potential role of the MYB–NFIB gene fusion, a recurrent chromosomal rearrangement in salivary gland ACC.

We identified a shared spatial localization among all MYB-expressing clusters in our well-structured analysis. While the molecular basis supporting this observation may not be obscure, it remains critical to validate shared transcriptional profiles, convergent oncogenic programming, and the implications of a common lineage origin from a singular transformed progenitor population confirming the underlying clonal evolution. These findings have important therapeutic relevance and play a pertinent role in advancing our understanding of how cellular states are modulated by tissue-specific spatial contexts and represents a crucial, yet underexplored, dimension in the study of this salivary gland neoplasm. We observed a pronounced upregulation of genes implicated in the disruption of cell-cell adhesion, extracellular matrix remodeling, and intermediate filament reorganization, which is evidenced by elevated expression of MMP7, FBLN, COL1A1, and POSTN. Additionally, a dysregulated suppression of canonical Wnt signaling was identified within the mildly MYB-expressing regions of the tissue architecture. This aberration is likely attributable to the intra-tumoral complexity characteristic of SGACC and the presence of a P-EMT phenotype. The P-EMT signature appears to be associated with the reactive stromal compartment, which is highly prominent in the spatial transcriptomic data and mirrors patterns observed in non-ACC malignancies.

Our study reinforces the classification of SGACC as an epithelial-derived malignancy, consistent with the fact that epithelial cancers comprise approximately 90% of all human tumors. While hallmark features of EMT were prominently observed in our analysis, an additional layer of evidence supporting the epithelial origin of the tumor lies in the marked upregulation of keratin expression within the non-MYB-expressing peripheral regions of the tumor architecture. We infer an epithelial progenitor origin for the root cell population based on the observed attenuation of keratin regulation, suggestive of a phenotypic shift toward vimentin-type intermediate filaments which is implicated in EMT programing^115^. This finding is further supported by the co-existence of epithelial and mesenchymal phenotypes, a feature also documented in carcinosarcomas, rare biphasic tumors arising in tissues such as the lung and uterus, where both lineages originate from a common epithelial progenitor^55^.

We identified a population of cancer-associated fibroblasts (CAFs), marked by the expression of genes such as COL1A1, COL3A1, COL11A1(Fig. 2D), and ACTA2, which are known to contribute to the activation of EMT programing^116,117^. This observation aligns with previous studies demonstrating the role of CAFs in promoting EMT, such as in co-culture models of PC-3 human prostate carcinoma cells, where CAF-mediated the secretion of matrix metalloproteinases (MMPs) such as MMP1, MMP2, MMP7, MMP9, and MMP11 drives EMT induction^118^ (Fig. 2D). These findings underscore the pivotal role of the TME, which is predominantly composed of extracellular matrix (ECM) components and non-transformed stromal cells, in fostering tumor progression through dynamic tumor-stroma interactions. In our study, such interactions were reflected by the exaggerated presence of fibroblastic and endothelial cells and the emergence of aberrant phenotypes, likely shaped by the reciprocal signaling between tumor cells and surrounding stroma. Particularly, the transcriptional upregulation of hepatic stellate cell activation pathways within the epithelial progenitor populations suggests a possible shared stromal remodeling mechanism, despite SGACC being a non-hepatic malignancy. While hepatic stellate cells in the liver typically function in vitamin A storage and ECM homeostasis, our findings point to a molecular convergence between CAF-associated pathways and stellate cell-like activation, potentially contributing to the pro-tumorigenic ECM reorganization observed in SGACC^119–122^.

Furthermore, analysis of the spatial transcriptomic profiling demonstrated a quantifiable infiltration of CD8⁺ T cells into the tumor core, along the fibroblastic tracks, indicating targeted immune cell migration toward tumoral niches. The analyzed SGACC sample, derived from a stage III–IV tumor, belonged to a patient who exhibited an extended survival period of over 204 months significantly surpassing the median survival duration typically reported for this malignancy.

In parallel, we observed evidence of plasma cell interactions within the TME, suggesting a role in modulating both tumor-promoting inflammation and immune surveillance. These interactions point to either an active or potentially dysregulated humoral immune response, as indicated by the upregulated expression of genes such as *ITGAX*, *IGLC2*, and *IGKC*. Such expression patterns may reflect transient attenuation of tumor progression mediated by antibody-producing cells, reinforcing the importance of both innate and adaptive immune processes in shaping SGACC pathophysiology^79,123,124^. Moreover, the pronounced spatial localization of plasma cell (PC), as confirmed by spatial mapping, supports a tertiary lymphoid structure (TLS)-dependent organization within the TME, offering a compelling framework for immunotherapeutic interventions.

Our analysis identified the upregulation of key signaling pathways, including the PI3K-Akt and IL-17 pathways, both of which have been previously implicated in adenoid cystic carcinoma (ACC) of the lacrimal gland. This supports the notion that these pathways are associated with the histological subtype of ACC rather than the anatomical site of tumor origin^125^. Strikingly, upregulation of the PI3K-Akt pathway is frequently linked to the activation of transmembrane receptor proteins such as *ITGB4*^126,127^, which was also observed in our dataset. Additionally, this pathway has been associated with immune-related conditions, including primary immunodeficiency highlighted in our disease pathway analyses and is similarly reported to be upregulated in individuals with acquired immunodeficiency syndrome (AIDS). Interestingly, this convergence has therapeutic implications, as nelfinavir, an antiretroviral drug commonly used in AIDS treatment, is currently under clinical investigation for its potential efficacy against ACC ^128^.

## 5.0 Conclusion

Our findings underscore the importance of identifying novel regulators of EMT for the development of more targeted and effective therapies for SGACC. Overall, our study presented the spatial integrity of the SGACC tumor landscape, enabling high-resolution visualization of gene expression patterns, cellular interactions within the TME, lineage tracing of tumor niches, and integration of spatial transcriptomic data with histological features corroborated by pathologist annotation collectively advancing our capacity to characterize and interpret non-dissociable tissue architectures that have not been reported until now.

## DECLARATIONS

### Ethics approval and consent to participate

De-identified adenoid cystic carcinoma tumor samples were obtained from several institutions: the Department of Otorhinolaryngology and Maxillofacial Surgery, Zealand University Hospital; the Department of Otorhinolaryngology, Head and Neck Surgery and Audiology, Rigshospitalet; the Department of Pathology, Rigshospitalet, University of Copenhagen; and the Department of Ophthalmology, Rigshospitalet-Glostrup, University of Copenhagen, Copenhagen, Denmark. All samples were provided Formalin-Fixed and Paraffin-Embedded (FFPE) as 5-micron sections baked onto glass slides. Salivary gland samples with survival information had at least 5-year follow-up. All samples were collected in accordance with the principle of the Declaration of Helsinki and with Institutional Review Board-approved protocols: Danish Regional Ethics Committee (H-6-2014-086) and the Danish Data Protection Agency (Journal no. REG-94-2014).

### Consent for publication

Informed consent was obtained from all subjects involved in the study.

## Availability of data and materials

The h5ad files will be made available.

## Author’s contribution

IE and GB conducted the research, developed various components of the software used in this study, and analyzed the data. IE drafted the manuscript, and GB contributed to manuscript editing. KH collected the data. KJB assisted in editing the manuscript and provided guidance regarding ACC tumor biology. ELB performed pathology tissue annotations and participated in manuscript revisions. SAN provided access to the ACC samples. JSE conceived the project, contributed to data analysis, and edited the manuscript.

## Acknowledgement

This research was partially supported by UNM Comprehensive Cancer Center Support Grant NCI P30CA118100. The authors wish to thank the Purdue University Rosen Center for Advanced Computing for providing Anvil computational resources funded by the National Science Foundation (Grant Number: 2005632). Access to computing resources was provided through the Advanced Cyberinfrastructure Coordination Ecosystem: Services & Support (ACCESS) program.

