## Supplementary Data for "Gene-Expression Programs in Salivary Gland Adenoid Cystic Carcinoma Analyzed Using Single-Cell and Spatial Transcriptomics"

### Integrative Multi-Omics Reveals Gene-Expression Programs in Salivary Gland Adenoid Cystic Carcinoma through Single-Cell and Spatial Transcriptomics

Ifeoma Ebinumoliseh, Gopikrishnan Bijukumar, Kendall Hoff, Kathryn Brayer, Elaine Bearer, Scott Ness, Jeremy Edwards

SUPPORTING INFORMATION:

**1A**

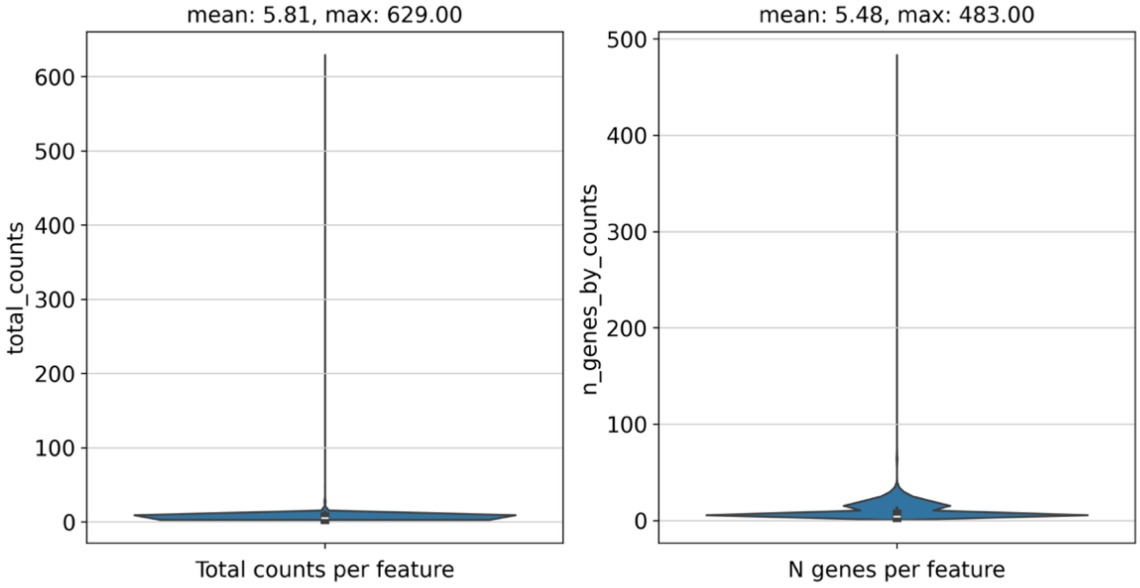


**1B**

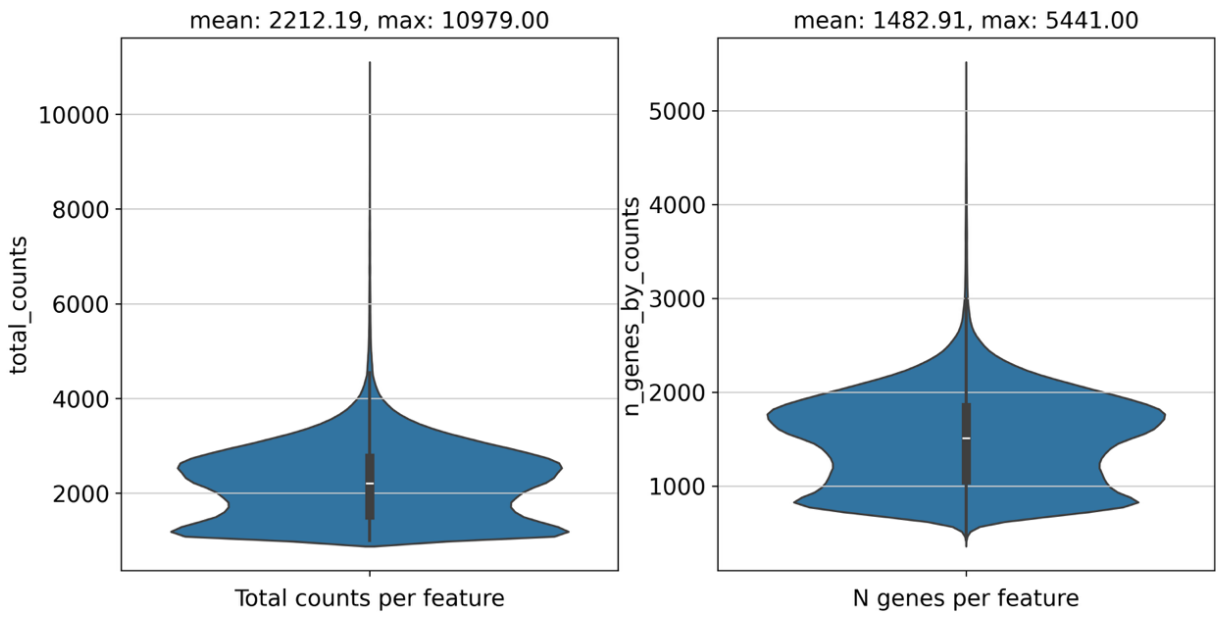


**Figure. S1A and B.** Preprocessing Quality control metrics of the spatial transcriptomics query dataset. (A) Total counts per feature and N-genes per feature before binning. (B) Total counts per feature and N-genes per feature after binning

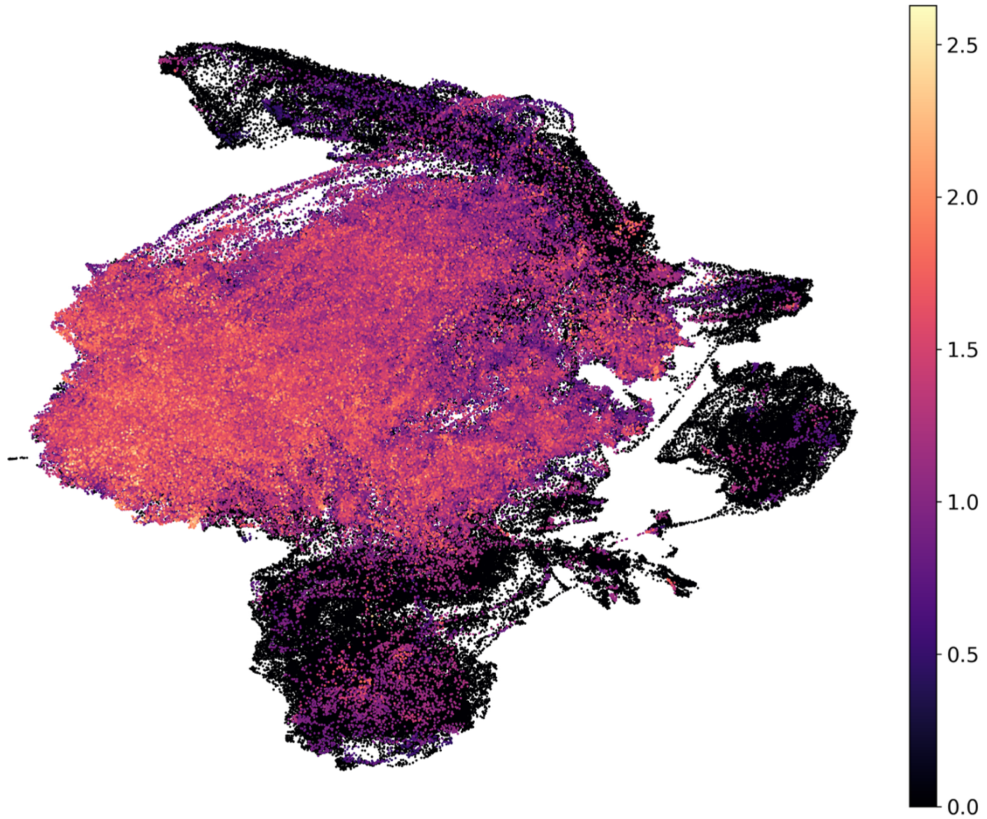


**Figure S2**. MYB Expression in UMAP of Spatial Data

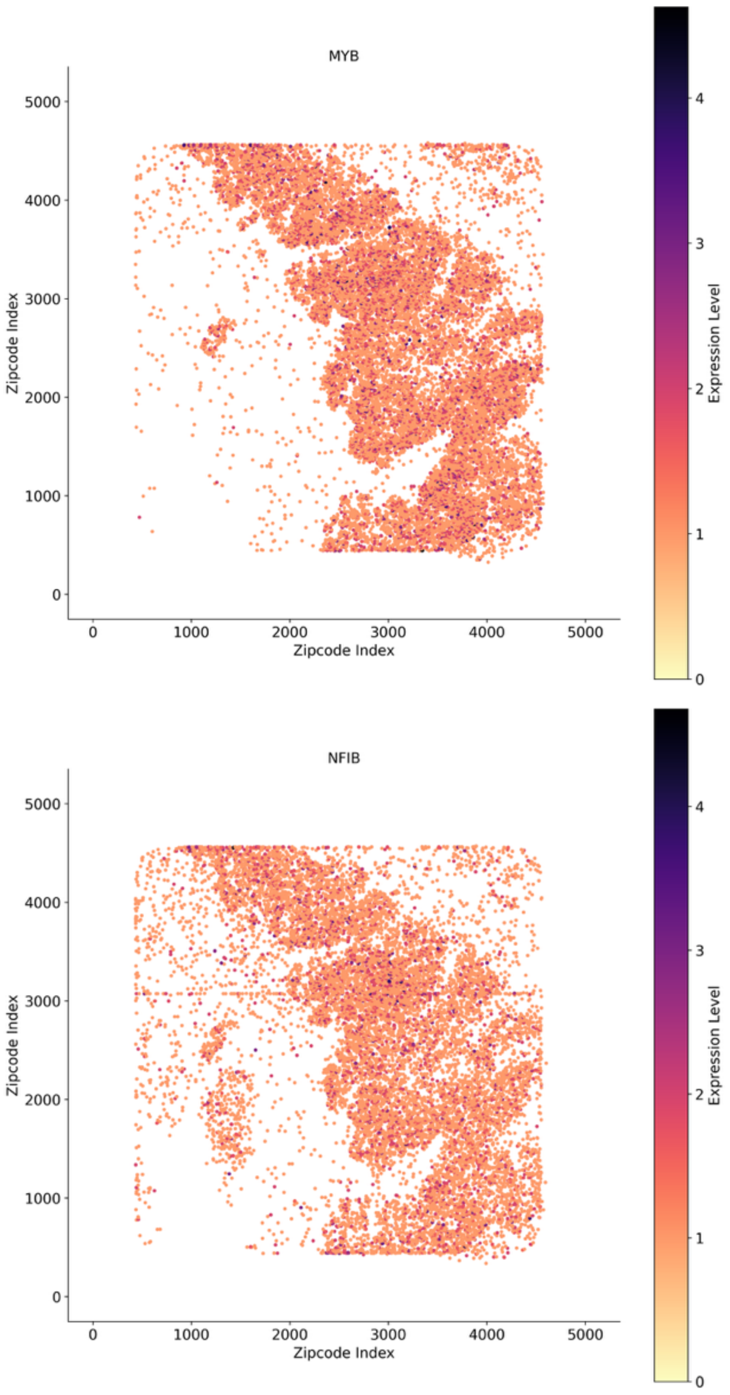


**Figure S3.** Mapped Zip Codes Showing MYB and NFIB Expression in Tissue Architecture

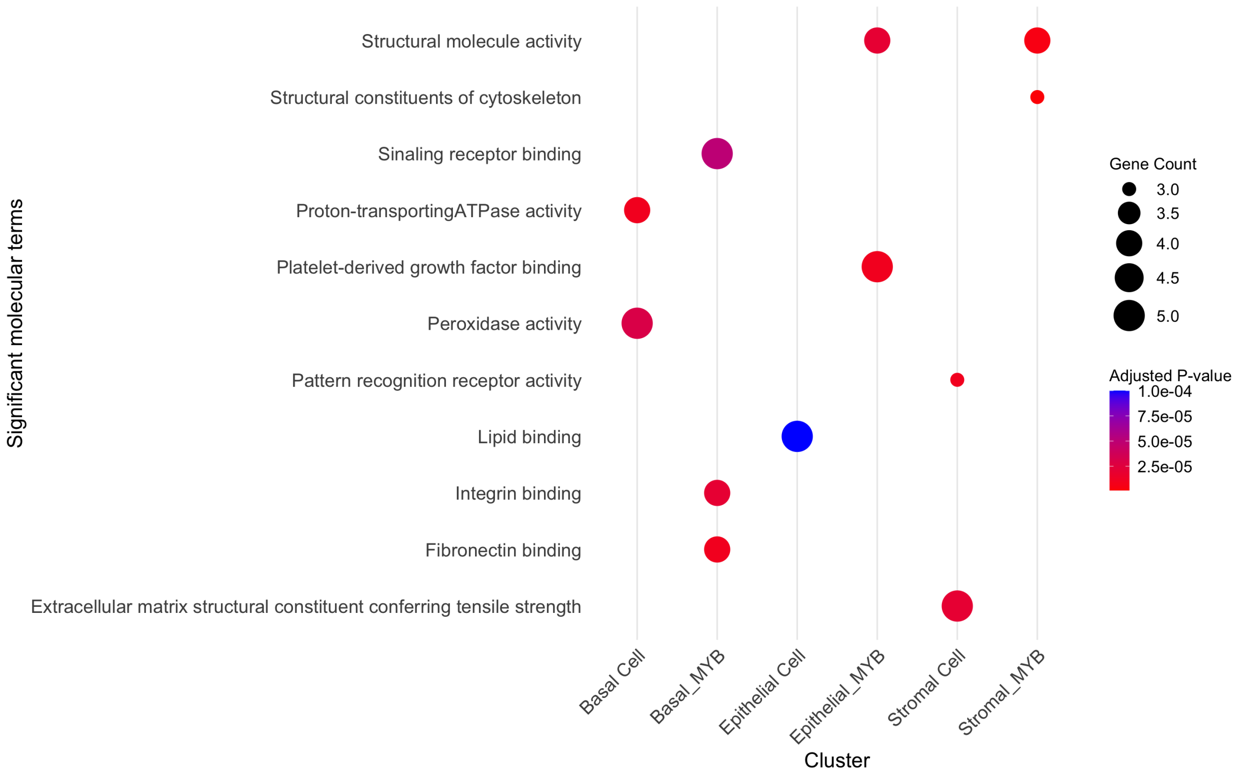


**Figure S4.** Pathway Enrichment of MYB and Non-MYB Expressing Stromal, Epithelial and Basal Clusters - Significant molecular terms

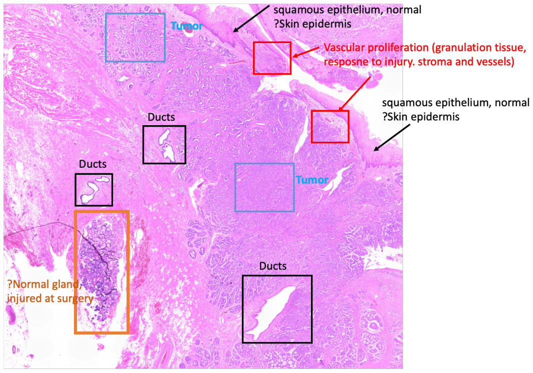


**Figure S5.** Pathology Annotation of Tissue

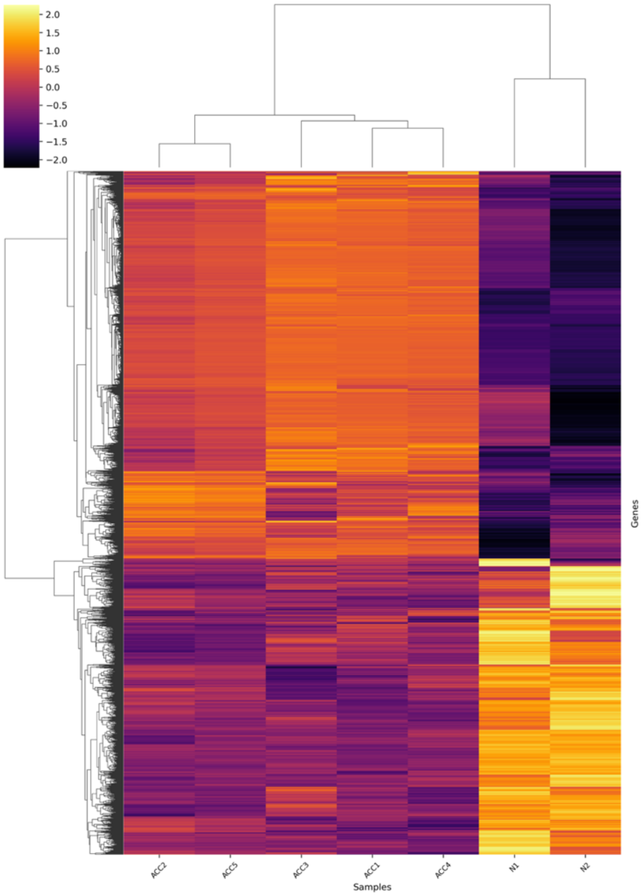


**Figure S6**:Heatmap of top 8,000 differentially expressed genes of normal control (N1 and N2) vs SGACC (ACC1 -ACC5) samples (n = 7)

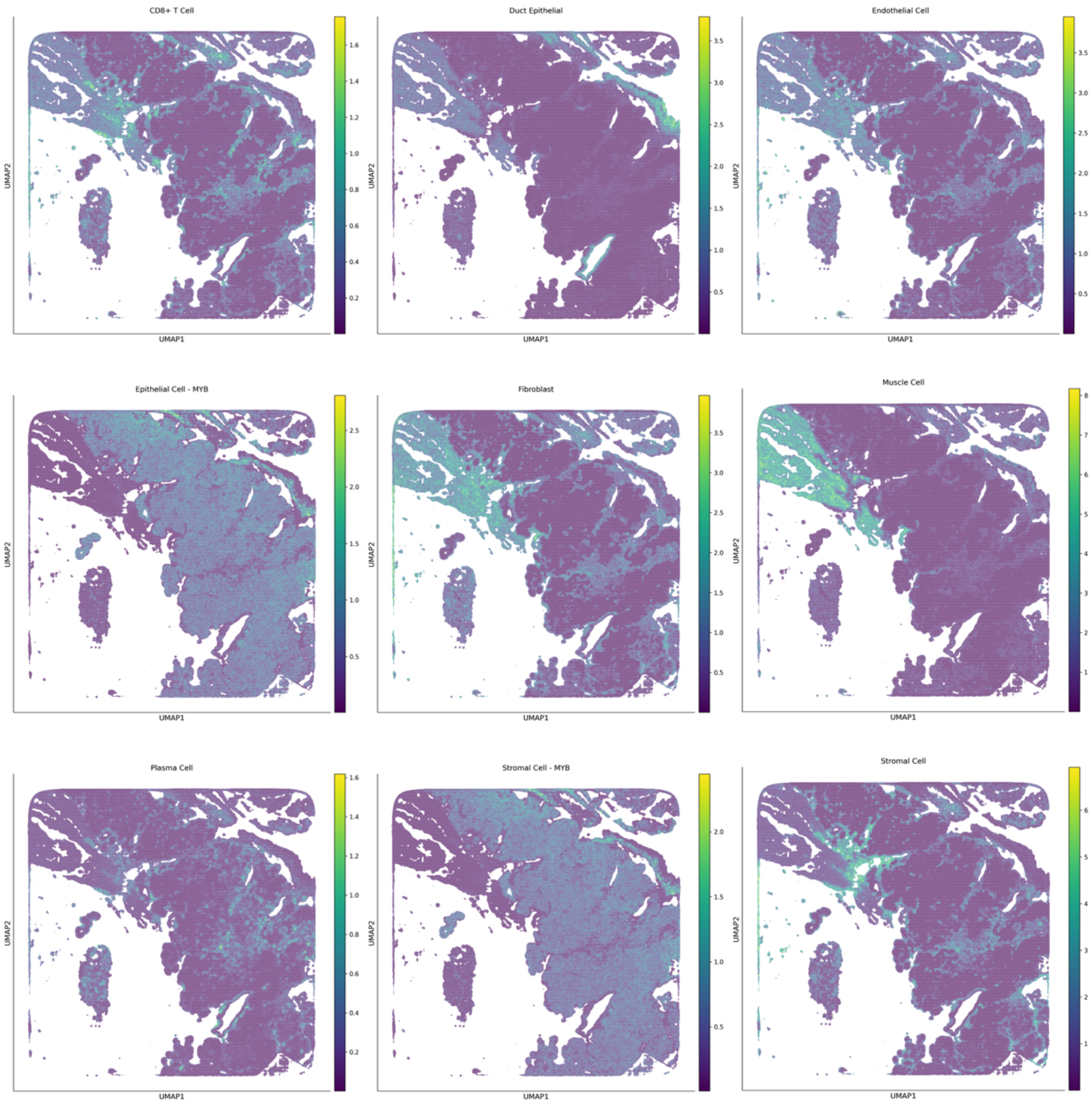


**Figure S7.** Spatial Localization of Individual Cells Within the Tissue Architecture

#### Table S1:

#### Marker genes for each cell types

| Epithelial Cell-MYB | Basal Cell-MYB | CD8+ T cell | Duct Epithelial Cell | Endothelial Cell | Epithelial Cell | Fibroblast Cell | Muscle Cell | Plasma Cell | Stromal Cell | Stromal Cell-MYB |
|---|---|---|---|---|---|---|---|---|---|---|
| **AZGP1** | KRT5 | GZMH | MUC5B | MGP | KRT13 | COL1A1 | TNNT3 | IGKC | CLEC3B | ALDH1A3 |
| **ITGB4** | KRT17 | KLRG1 | PIGR | EGR3 | SPRR2A | COL3A1 | CKM | IGHA2 | COL1A1 | EGR3 |
| | | | BPIFB2 | | SBSN | COL1A2 | MB | IGLC2 | HBA2 | NR4A1 |
| | | | | | LY6D | | TTN | JCHAIN | | |
| | | | | | | | STAC3 | | | |
